# Predictive pursuit emerges in high-dimensional recurrent neural networks

**DOI:** 10.64898/2026.04.23.720457

**Authors:** William T. Redman, Fatih Dinc, Xiaoxiao Lin, May G. Chan, Andrew S. Alexander

## Abstract

Tracking dynamic moving objects in the external world is ethologically important for many organisms. Recent experiments have examined neural dynamics supporting such behaviors by employing visually-guided pursuit in freely moving rodents, yet computational principles underlying this cognitive process are not well understood. To address this, we developed a recurrent neural network model for examining the predictive behaviors and computations that emerge during pursuit. We demonstrate that the model generates internal predictions of the target’s future locations, with anticipatory behaviors increasing with exposure to stereotyped trajectories of the target. These internal predictions can be used by the model to pursue a target in a complex environment, and the model’s emergent strategy is aligned with behavior when tested in rodents. In investigating the computations that underlie the model’s ability to perform predictive pursuit, we found units sensitive to the position of the target relative to the artificial agent, a representation analogous to egocentric target neurons observed in animals performing pursuit tasks. Ablating these units significantly reduced model performance, establishing a causal role of this functional response type in efficient pursuit. Given the complexity of the task and agent behavior, we hypothesized that RNN models may use high-dimensional neural codes to support predictive pursuit. To test this, we trained models of varying rank and found that anticipatory behavior emerged only when the rank was sufficiently high, despite strong pursuit performance in lower rank models. All RNNs encoded the egocentric location of the target, whereas allocentric self and target locations emerged only in high-dimensional networks. Overall, our results suggest that, unlike commonly studied vision, motor, or memory tasks, predictive pursuit emerges in high-dimensional networks with sufficient resources.

## Introduction

Pursuit is a ubiquitous behavior in biology, with a wide a range of organisms chasing others for predation and reproduction. In cases where the pursuer is faster than its target, moving to minimize the relative distance between the two can be an effective strategy for capture. However, when the target can move at higher speeds than the pursuer or is more agile, effective pursuit requires utilizing dynamic information about the target to make predictions of where it is headed. The complex way in which environmental affordances (e.g., boundaries, burrows), movement statistics (e.g., range of typical speeds), and stereotyped actions (e.g., common routes, zigzagging, slowing and accelerating) inform this prediction makes pursuit a rich paradigm for studying the neural computations underlying model-based inference and predictive planning [1].

Evidence of the ability to predictively pursue others has been found in dragonflies [2], snakes [3], bats [4–6], rats [7], and non-human primates [8, 9], demonstrating the widespread use of this behavior. For some organisms (e.g., snakes), the neural circuitry enabling this predictivity may be evolutionarily hard-wired. However, for other organisms (e.g., rats and non-human primates), predictive pursuit can emerge with learning [7–9]. As planning and inference drive many intelligent behaviors, a central challenge in neuroscience is to identify neural computations that enable prediction.

To address this open question, we develop a recurrent neural network (RNN) model for examining the behavior and computations that emerge during continuous pursuit of a moving target. RNN models are powerful tools for normatively studying the computations that underlie cognitive processes [10–17] and have been used to study observed neural computational properties in spatial navigation [18–22] and decision making [23–26] tasks. In addition, RNNs enable manipulations not easily possible in systems neuroscience experiments, such as ablating specific functional classes of neurons [27, 28] and changing the inherent rank of the network’s connectivity [29–38] (i.e., the number of degrees of freedom with which the recurrent units of the RNN are wired together).

Here, we interrogate RNNs trained on pursuit tasks. We find that RNNs can accurately intercept targets, and learn to do so using strategies that suggest the development and deployment of an internal model. Specifically, we find that RNNs can anticipate future locations of the target, generating pursuit trajectories that are significantly more predictive than those exhibited by a non-predictive control model. Model predictive behaviors can be maintained even when information about the target is intermittently masked, supporting an internal model of target behavior during pursuit. RNNs make experimentally testable predictions about how a predictive biological agent behaves during novel target pursuit tasks, which we validate against newly collected rodent behavioral data. Beyond behavior, we find representations in the RNNs that resemble neural egocentric receptive fields reported in target-chasing rats [39, 40], and lesions of these units disrupt model performance. Collectively, these results demonstrate that the RNNs learn complex, predictive pursuit strategies that closely parallel rodent behavior at both the single-unit and behavioral levels.

To analyze how the RNN learns predictive pursuit strategies, we perform a mechanistic study of predictive pursuit in networks with constrained connectivity (i.e., low-rank RNNs). Earlier studies have primarily focused on visual, motor, or memory tasks that are often solvable via low-dimensional networks with limited computational resources [13, 30–34, 36]. When trained on such tasks, even when networks are not explicitly constrained, RNNs tend to converge to low-rank solutions after training [32, 41, 42]. This observation is generally attributed to the inherent low-dimensionality of the behavior they model [11, 24, 43]. Predictive pursuit, by contrast, requires modeling the environment and the target, a complex demand that we hypothesize would require high-dimensional network connectivity. To test this, we train RNNs of varying rank and examine their pursuit trajectories. We find that anticipatory behavior emerges only in RNNs that are high-dimensional, even though lower rank RNNs solve the task with high accuracy. Examining the learned representations of the models, we find that all RNNs learn to represent the egocentric location of the target. However, only higher rank RNNs learn to represent allocentric self and target locations, in which allocentric representations presumably support anticipatory trajectory planning. This suggests that complex, ethologically relevant behaviors may require high-dimensional network solutions, which are inherently harder to interpret than the low-dimensional solutions that suffice for idealized tasks [24, 31]. As the field moves toward studying neural computation in naturalistic settings [44, 45], our work constitutes a mechanistic step in this direction, providing a tractable RNN framework for studying the rich computational structure that underlies predictive pursuit.

## Results

### RNN models learn to pursue moving target

We develop an RNN model that learns to steer an agent (“RNN-agent”) in pursuit of a “target-agent”. The RNN takes as inputs the initial positions of both agents and the instantaneous velocity of the target-agent, and outputs its own velocity (Fig. 1A; see Appendix A for details). The RNN is trained to move the RNN-agent towards the target-agent, so that the distance between the two on the last time-step of the trial is minimized. This is done to mimic “capturing” the target, but in a supervised learning setting. Each trial begins with the RNN-agent and target-agent initialized at distinct locations and lasts for *T* time-steps. Importantly, except for the starting positions, (*x*_RNN_(0), *y*_RNN_(0)) and (*x*_target_(0), *x*_target_(0)), the RNN is not given any ground truth spatial information. Thus, to solve the pursuit task, the RNN must integrate the dynamic information of both agents [46].

**Figure 1:**
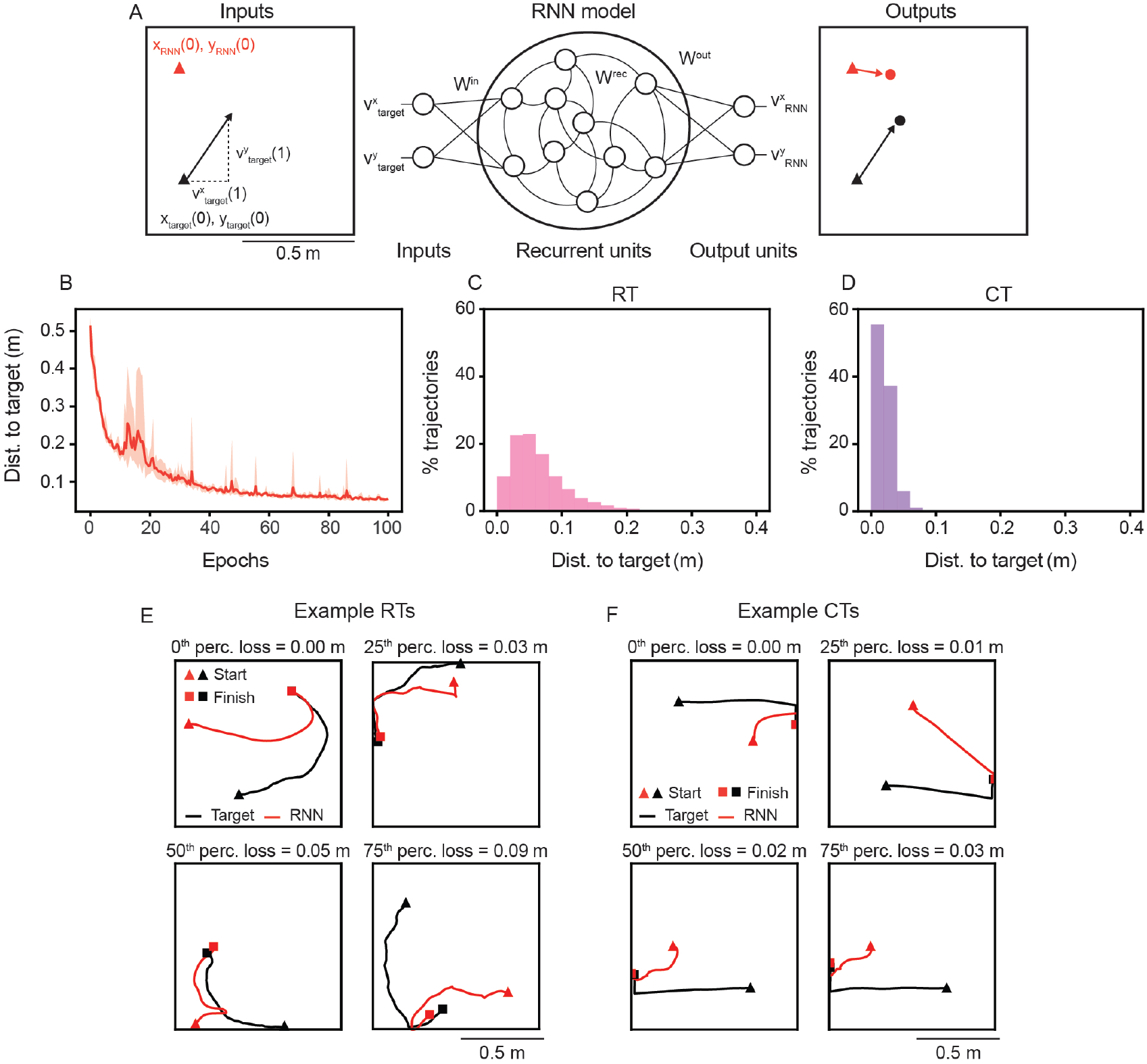
RNNs learn to pursue a moving target. (A) Schematic of the RNN model architecture. (B) Distance to target-agent at the end of the trial, which is used as the training loss, as a function of training epochs. Solid line is mean and shaded area is minimum and maximum of five independently trained networks. (C)–(D) Distribution of distance to target at the end of the trial across all 5000 sampled pseudo-random trajectories (RTs – 1000 per network) and characteristic trajectories (CTs – 1000 per network). (E)–(F) Example RT and CT trials. Trials were chosen as those closest to the 0^th^, 25^th^, 50^th^, and 75^th^ percentiles of the distance to target-agent distribution, respectively.

We design two kinds of trials, in which the target-agent exhibits one of two behaviors: random trajectories (RTs) and characteristic trajectories (CTs). During RT trials, the target-agent pseudo-randomly samples movement directions and speeds at each time-step, with a bias for smooth trajectories and avoidance of walls [19] (Fig. 1E, black lines). This makes each RT trial unique. During CT trials, the target-agent starts from one of four locations, heads towards the opposite wall, and then moves along the wall towards the middle of the environment (Fig. 1F, black lines). While the exact speeds and movement directions sampled vary on each trial, leading to slight differences, the overall structure of the target-agent’s trajectory is the same across CT trials (see Appendix B, for details). These two types of trials are similar to those used in recent rodent pursuit experiments [7].

In this work, we train our RNN model on a mixture of both RTs and CTs (75% RTs and 25% CTs for most of our analysis; this approximately matches rodent pursuit experiments [7]). We find that the RNN learns to pursue the moving target-agent with high accuracy (Fig. 1B), achieving a median distance from the end location of the target-agent of approximately 0.05 m on RTs and 0.02 m on CTs (Fig. 1C, D; arena is of size 1 m *×* 1 m). Visual inspection of individual trials confirms that the RNN develops complex pursuit behaviors (Fig. 1E, F), including sharp turns (Fig. 1E, lower left) and paths that appear to anticipate the future trajectory of the target (Fig. 1F, top right). Below, we first analyze the neural representations learned by the RNN that enables pursuit and then quantify the predictive behaviors.

### RNN models leverage egocentric representations for pursuit

Electrophysiological recordings in rats pursuing a dynamic target (in trials similar to our RTs) have identified neurons in posterior parietal cortex (PPC) and retrosplenial cortex (RSC) that demonstrate tuning to the location and orientation of the target relative to the self [7, 40]. These egocentric target cells (ETCs) were hypothesized to play an important role in pursuit during RTs, although causally demonstrating this in freely behaving animals is challenging. Given this putative role of ETCs in pursuit, we asked whether such a representation emerged at the level of individual units in the RNN. Plotting the average unit activations as functions of egocentric target position, we find units whose activations are selectively increased when the target-agent is at specific distances and orientations relative to the RNN-agent (Fig. 2A, top row). The egocentric target tuning of these units becomes especially clear when comparing them to units that exhibit no such selectivity (Fig. 2A, bottom row).

**Figure 2:**
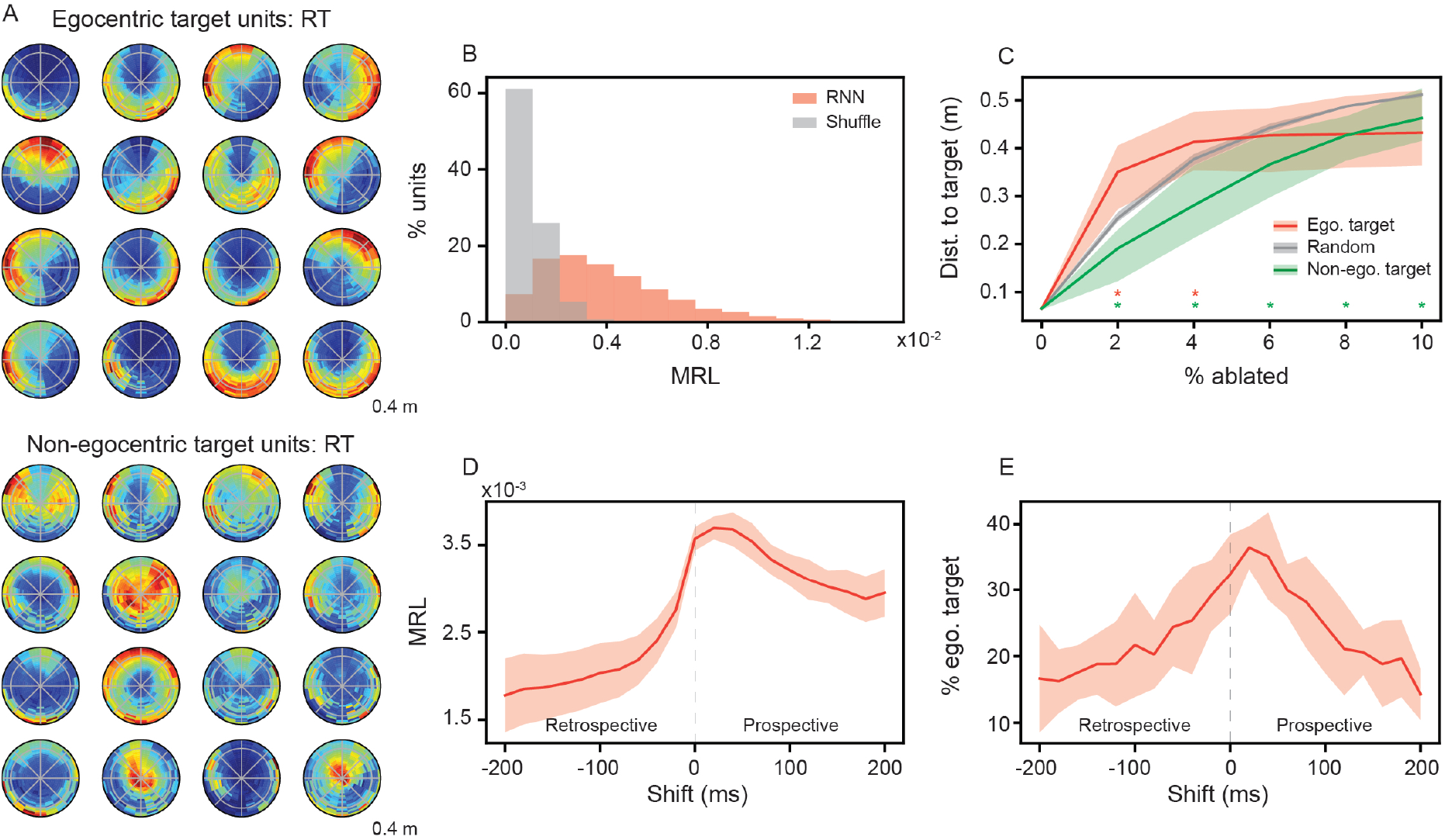
Egocentric target representations are causally linked to pursuit in RNNs. (A) Example ratemaps of units with strong egocentric target tuning (top row) and example ratemaps of units with weak egocentric target tuning (bottom row). (B) Distribution of mean resultant length (MRL) for RNN units (red) and shuffle control (gray). (C) Performance of RNNs when ablating units with the highest MRL (red), the lowest MRL (green), or by random selection (gray). Solid line is mean and shaded area is minimum and maximum of five independently trained networks. Stars denote ablation percentages where a significant difference to the random unit ablations exists (*p* < 0.05 using two-sample Kolmogorov-Smirnov test) (D) Average MRL, across all units, when shifting the target-agent’s position either prospectively or retrospectively. (E) The percent of all units classified as ETUs, when shifting the target-agent’s position either prospectively or retrospectively. Solid line is mean and shaded area is minimum and maximum of five independently trained networks.

A standard method for quantifying the strength of egocentric representations is through the computation of the associated mean resultant length (MRL) [7, 39]. This metric is independent of the distance between two agents and quantifies how strongly a neuron’s activity is concentrated around a preferred egocentric direction to the target (see Appendix C for details). We compute the MRL for all units in the RNN models and compare them to a shuffle control, where the activations are randomly shuffled across time points for each unit (Fig. 2B; see Appendix C for details). We classify units as egocentric target units (ETUs) if their MRL is greater than the 99^th^ percentile of the shuffled distribution and their MRL values are consistent across two splits of trials. Overall, we find 36% of all recurrent units are ETUs (range: 29%-42% across five independently trained RNNs), which is consistent with proportions observed in neural recordings [7].

To understand whether this emergent egocentric target representation is used by the RNN to perform pursuit, we order ETUs by their MRL and remove (or “ablate”) an increasing number of them from the model. We find that the RNN is sensitive to the loss of ETUs and performance quickly deteriorates significantly more than when removing randomly selected units (Fig. 2C; compare red and gray curves). In addition, we find that removing units with the smallest MRL values impacts model pursuit performance significantly less than when removing randomly selected units (Fig. 2C; compare green and gray curves). Together, these observations establish a causal role for ETU representations in successful pursuit in RNNs.

To further characterize the nature of these representations, we next examine whether ETU activity reflects past, current, or future target states. Specifically, we shift the egocentric target coordinates forward (“prospective”) or backwards (“retrospective”) in time, relative to the unit activations. Recomputing the egocentric target ratemaps on the shifted data, we find that the average MRL exhibits strong decreases for retrospective shifts, but not prospective shifts (Fig. 2D). In addition, the percent of units classified as ETUs peaks at a slight prospective shift (Fig. 2E). This suggests that the emergent egocentric target representations are not merely products of encoding the past relative positions of the target-agent, but instead skewed towards encoding the current and future locations of the target-agent. This greater prospective, as opposed to retrospective, coding further supports ETU’s functional utility in the pursuit task.

### RNN models learn to conduct predictive pursuit

While the RNNs are able to successfully pursue the target-agent, this does not imply that they are doing it in a *predictive* manner as observed in rodents or other species [2, 6–8]. For instance, the RNN-agent could simply learn to minimize the distance with the target-agent at every time step (i.e., pure pursuit). Since RNNs are scored based on the final distance between two agents, this strategy could still represent one minimum of the loss function. However, our analysis in Fig. 2D, E reveals prospective coding in ETUs, a result suggestive of the RNN models’ capability of performing predictive pursuit. We therefore quantify and test the capabilities of the RNNs to conduct pursuit with predictive strategies.

We compare the pursuit trajectories generated by the RNNs to a non-predictive control model, which moves in the direction that minimizes its distance to the current position of the target-agent (Fig. 3A). This control model is implemented by a set of equations (see Appendix D for details). The control model uses the same instantaneous speed as that outputted by the RNN model (as opposed to the maximum possible speed), meaning it does not perform an optimal greedy pursuit, but rather a realistic approximation of *reactive pursuit*. By construction, this reactive (non-predictive) model ignores all the dynamical history of the target-agent that is inputted to the RNN, as well as any information about the boundaries of the environment, which may act to constrain the future behavior of the target-agent. Hence, if the pursuit strategy used by the RNN seeks to minimize instantaneous distance with the target-agent, then the non-predictive model should have the same trajectory as the RNN-agent. However, deviations from the non-predictive control trajectory alone do not necessarily imply prediction, as they could also reflect a poorer pursuit strategy. We therefore use the non-predictive control as a baseline for reactive pursuit, and quantify predictive behavior using geometric metrics derived from the resulting trajectories (described below).

**Figure 3:**
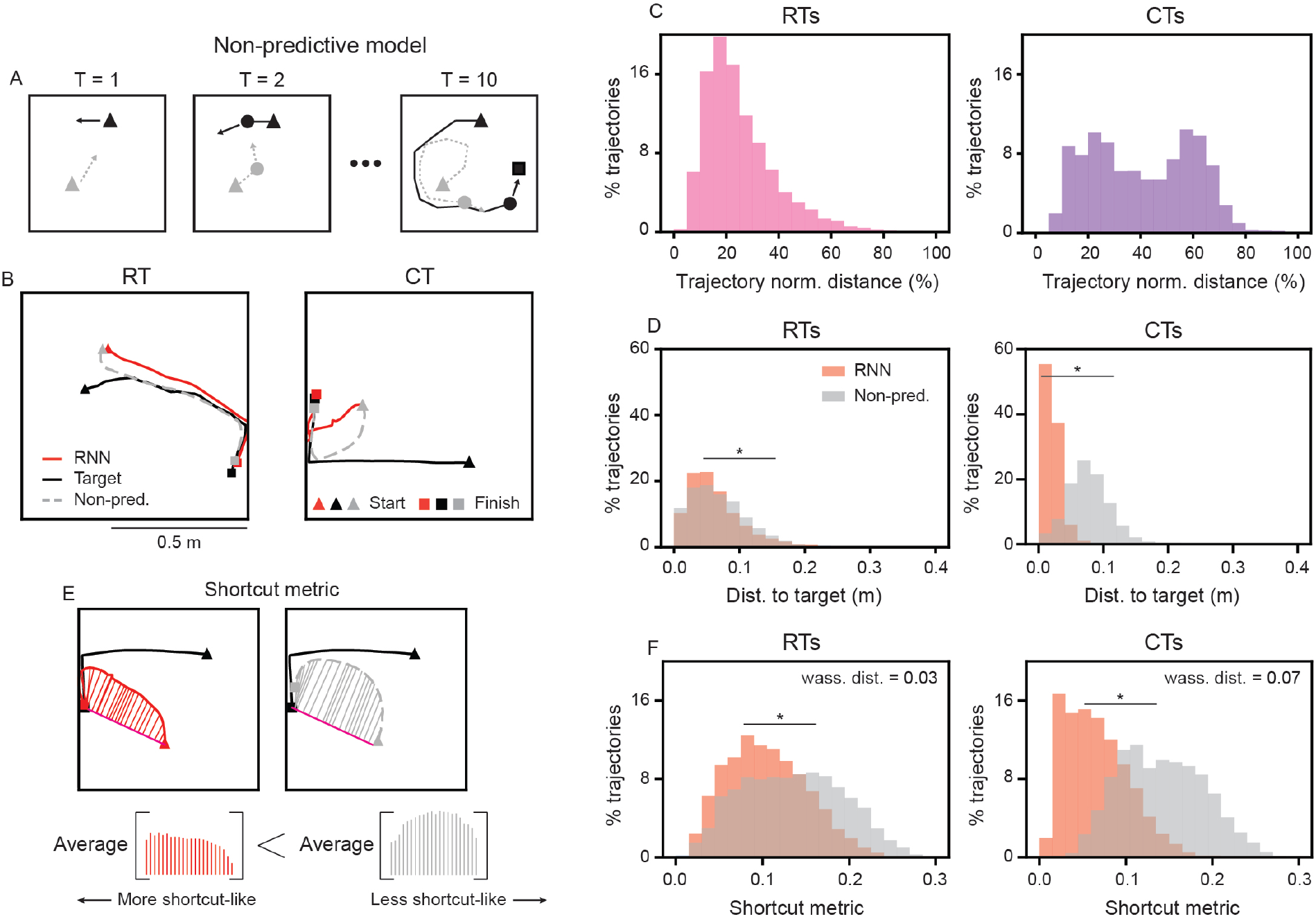
RNNs develop predictive pursuit behavior. (A) Schematic illustration of how the non-predictive control model updates its position. (B) Example RT and CT trials comparing the trajectories of the RNN (red solid line) and non-predictive control model (gray dashed line). (C) Distribution of normalized distance between RNN and non-predictive control model trajectories on RTs and CTs. (D) Distribution of distance to target-agent at the end of the trial for RNN and non-predictive control models on RTs and CTs. (E) Schematic illustration of the shortcut metric. Each model’s trajectory is compared to the direct path (pink line) from the start location of the RNN/non-predictive control model and the final location of the target-agent. (F) Distribution of shortcut metric for the RNN and non-predictive control models on RTs and CTs. Star denotes significant difference (*p <* 0.05 using two-sample Kolmogorov-Smirnov test).

Visually inspecting pursuit trajectories of both models, we find that the RNN performs pursuit along trajectories that differ from those used by the non-predictive model, on both RTs and CTs (Fig. 3B). Notably, the RNN and non-predictive control models’ trajectories differ on average by 22% of the total length traveled on RTs, and 40% of the total length traveled on CTs (Fig. 3C). These differences also manifest in differences in performance, as the RNN model outperforms the non-predictive control model, particularly on CTs (Fig. 3D; median distance to end location of target-agent on RTs: RNN = 0.054 m vs. non-predictive control = 0.062 m; CTs: RNN = 0.018 m vs. non-predictive control = 0.077 m).

Next, we quantify the difference in predictivity between the RNN and non-predictive control models’ trajectories by defining a geometrically motivated “shortcut metric”. First, we define a reference trajectory given by the direct path between the starting location of the RNN-agent and the final location of the target-agent (i.e., the path an agent with perfect knowledge would take; Fig. 3E). For each model, we then compute the average distance between its trajectory and this reference path. This quantity serves as a measure of how “shortcut-like” a trajectory is (see Appendix D.1 for more details), as an agent with full knowledge of the target’s final position should in principle follow this path, yielding a lower bound of 0. As a baseline for non-shortcut-like trajectories, we compare against the non-predictive control model.

We find that, on both RTs and CTs, the RNN model’s trajectories have significantly smaller shortcut metric (i.e., are more shortcut-like) than the non-predictive model (Fig. 3F; two-sample Kolmogorov-Smirnov test between RNN and non-predictive control model shortcut metrics *D*(5000) = 0.23, *p <* 0.001 on RTs and *D*(5000) = 0.58, *p <* 0.001, on CTs). We measure the difference in shortcut metric distribution across trials using the Wasserstein distance, where greater distance corresponds to more distinct strategies between the RNN and non-predictive control models. We find that the Wasserstein distance between the shortcut metric distributions of the RNN and non-predictive models is more than doubled on CTs when compared to RTs (Fig. 3F; Wasserstein distance = 0.03 on RTs and 0.07 on CTs), demonstrating that the presence of repeated structure is more heavily exploited by the RNN model. Furthermore, the significant difference between shortcut metrics on RTs, albeit being weaker compared to CTs, suggests that the RNN utilizes repeated aspects of the target-agent’s pseudo-random trajectory (e.g., its bias away from walls, its bias for smooth paths) to generate more predictive trajectories than would be expected if it only followed the target-agent. Overall, these results provide evidence that the RNNs employ predictive strategies during pursuit, rather than relying solely on reactive distance minimization.

### RNN model maintains and updates internal representation of target

A hallmark of predictive computation is the maintenance, updating, and use of an internal model of the environment, which in turn shapes ongoing behavior and neural processing [47–50]. Beyond generating predictions about future states, an internal model is expected to be acquired through structured experience and to persist autonomously even when sensory input is removed. We have already provided evidence for the predictive nature of the RNNs through representational (Fig. 2) and behavioral (Fig. 3) signatures. Here, we provide two further lines of evidence. Namely, that the RNNs’ predictive behavior is shaped by increasing exposure to structured trials (Fig. S1) and that RNNs can perform predictive pursuit even when a fraction of the dynamical information about the target-agent is masked (Fig. 4).

**Figure 4:**
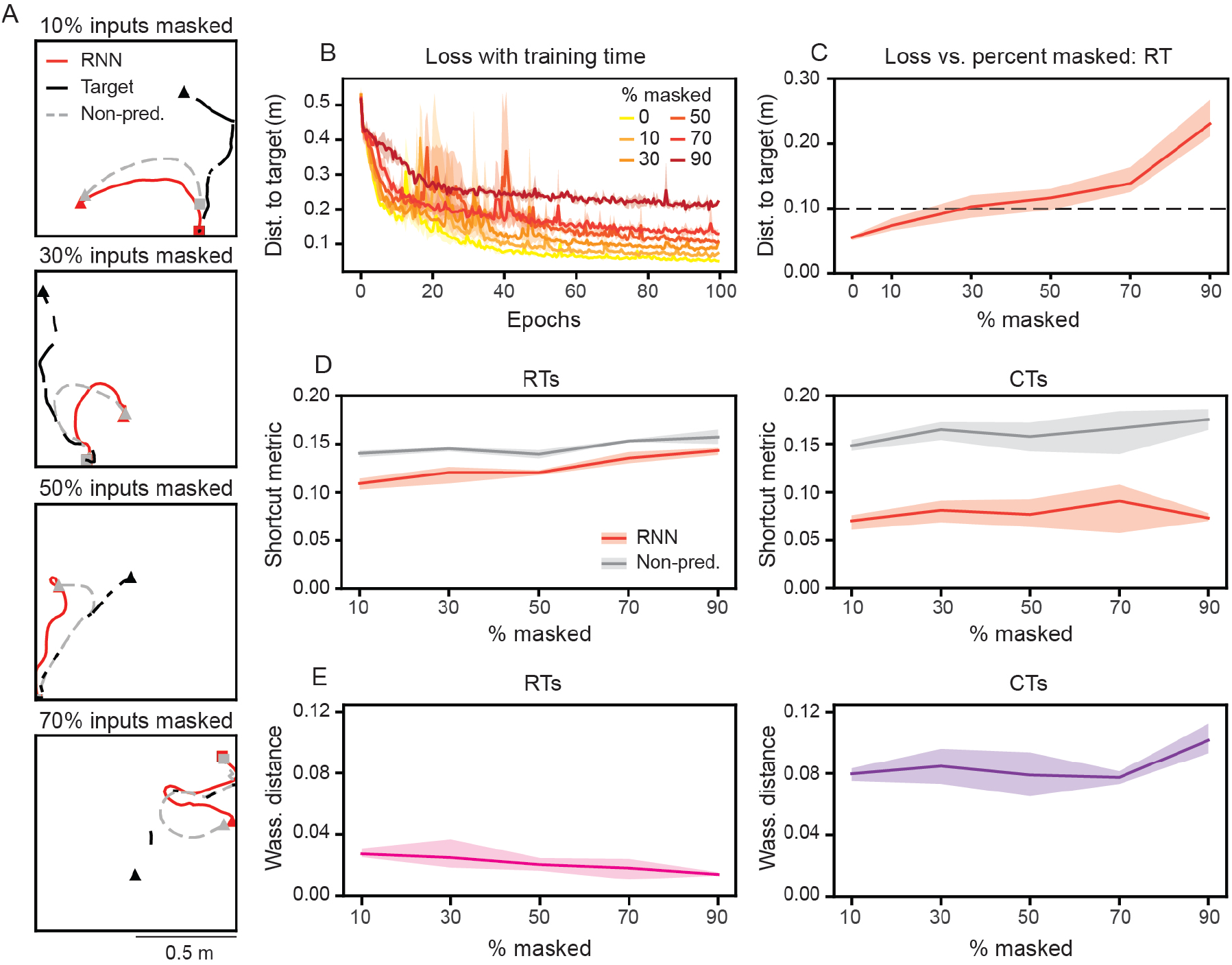
RNNs maintain an internal representation of the target-agent. (A) Example trajectories with 10%, 30%, 50%, and 70% of the RNNs’ inputs masked. Note that the non-predictive control model’s trajectory is computed assuming complete knowledge of the target-agent. (B) Distance to target-agent as a function of training epochs for RNNs trained with different amounts of input masking. (C) Distance to target-agent, as a function of input masking. Dashed black line denotes a distance to target-agent of 0.10 m. (D) Average shortcut metric for RTs and CTs, as a function of input masking. (E) Wasserstein distance between the distribution of RNN and non-predictive control model shortcut metrics for RTs and CTs, a sa function of input masking. (B)–(E) Solid line is mean and shaded area is minimum and maximum of three independently trained networks.

To start with, if the RNNs are indeed leveraging an internal representation of the target-agent to guide anticipatory trajectories during CT trials, then we would expect the strength of this behavior to be dependent on exposure to structured trials (i.e., CTs) during training. Thus, we hypothesize that this anticipatory behavior during CTs should be significantly reduced if the RNNs were trained only on RTs. We test this hypothesis by computing the average shortcut metric of RNNs trained on varying percentages of CT trials. We find that RNNs trained solely on RT trials have significantly larger shortcut metrics (i.e., are less shortcut-like) than RNN models whose training included at least 10% CT trials (Fig. S1A). This difference significantly increases as more CT trials are used in the training dataset, which we quantified by computing the Wasserstein distance between the distributions of shortcut metrics belonging to the RNN and non-predictive models (Fig. S1B; linear regression on Wasserstein distance as a function of CT % trained on; slope = 0.06, *R*^2^ = 0.73). Overall, as expected, RNN’s predictivity is enhanced with increased exposure to structure during training.

Next, if RNNs are leveraging an internal representation of the target-agent to plan their pursuit trajectories, then they should continue to perform well even when information of the target-agent is occluded. To test this, we train RNNs with varying percent of the inputs “masked out” (Fig. 4A). Here, we still provide the RNNs with the initial location information of both RNN-agent and target-agent, but provide the velocity of the target-agent only at a fraction of the time steps, inputting 0 otherwise. As the mask percent is increased, the RNNs receive fewer inputs (or equivalently, experience an increase in the temporal spacing between inputs). Unsurprisingly, we find that greater masking percentages lead to slower learning and worse overall performance (Fig. 4B). However, RNNs are still capable of conducting pursuit accurately (Fig. 4A, D). For instance, even when information about the target-agent is only input half the time, the RNNs can still get to within approximately 0.1 m of the target-agent on RT trials (Fig. 4C). Furthermore, the RNNs demonstrate evidence of predictive behavior (Fig. 4D, E). Masking the inputs more than 50% of the time leads to less difference between the RNN and non-predictive models’ trajectories on RT trials (Fig. 4D, E; left column), suggesting that large gaps in input leads the RNNs to learn a more reactive strategy on RT trials. In contrast, for CT trials, the RNNs maintain shortcut-like behavior, even as the masked percent increases (Fig. 4D, E; right column). This demonstrates that the repeated structure of the CTs can still be learned and exploited, even when instantaneous information about the target is withheld from the RNN models.

Taken together, RNNs show hallmark properties of systems with an internal model of their environment. Namely, their predictive behavior is grounded in representational and behavioral signatures, shaped by structured experience, and maintained autonomously in the absence of complete sensory input [2, 51, 52].

### Task and environmental determinants of predictive pursuit

We hypothesized that the predictive behavior learned by the RNNs was encouraged by the task objective, which penalized the end distance between the RNN-agent and target-agent. We tested this by changing the loss function such that the RNNs were trained to minimize the average distance between the target-agent and the RNN-agent at each time step. We reasoned that the need to minimize the distance at all times would incentivize a reactive pursuit strategy. As expected and shown in Fig. S2, the differences in behavior between the RNN and non-predictive controls (observed in Fig. 3) is weakened. In particular, when the RNNs are trained to minimize the mean distance between the RNN-agent and the target-agent across the trial, the learned trajectories are similar to their non-predictive controls on both RTs and CTs (Fig. S2; see Appendix E for more details). Thus, the predictivity (or conversely, the reactivity) of the RNN models can be controlled by the choice of loss function that defines task.

Next, to explore the environmental determinants of predictive pursuit, we train an additional set of networks on an environment with periodic boundaries. In this setting, the target-agent is allowed to pass through the boundaries and appear on the opposite side of the environment (Fig. 5A; black lines in left column). This addition makes the RT trials significantly harder (median distance to the end location of the target-agent in RT trials = 0.34 m), as the target-agent can quickly cover large amounts of distance through the connectedness of the boundaries. However, the RNN is still able to perform with high accuracy on CTs (Fig. 5A, left column; median distance to end location of target-agent in CT trials = 0.04 m). In these trials, the RNN generates trajectories that move towards the side of the environment where the target-agent will appear, as opposed to following the target-agent along its movement direction (Fig. 5A; left column, compare red and black lines). This predictive behavior is accompanied by a waiting strategy, in which the RNN-agent moves towards the boundary where the target-agent will appear and then waits (Fig. 5B; black triangles, see Fig. 5C for trial averages).

**Figure 5:**
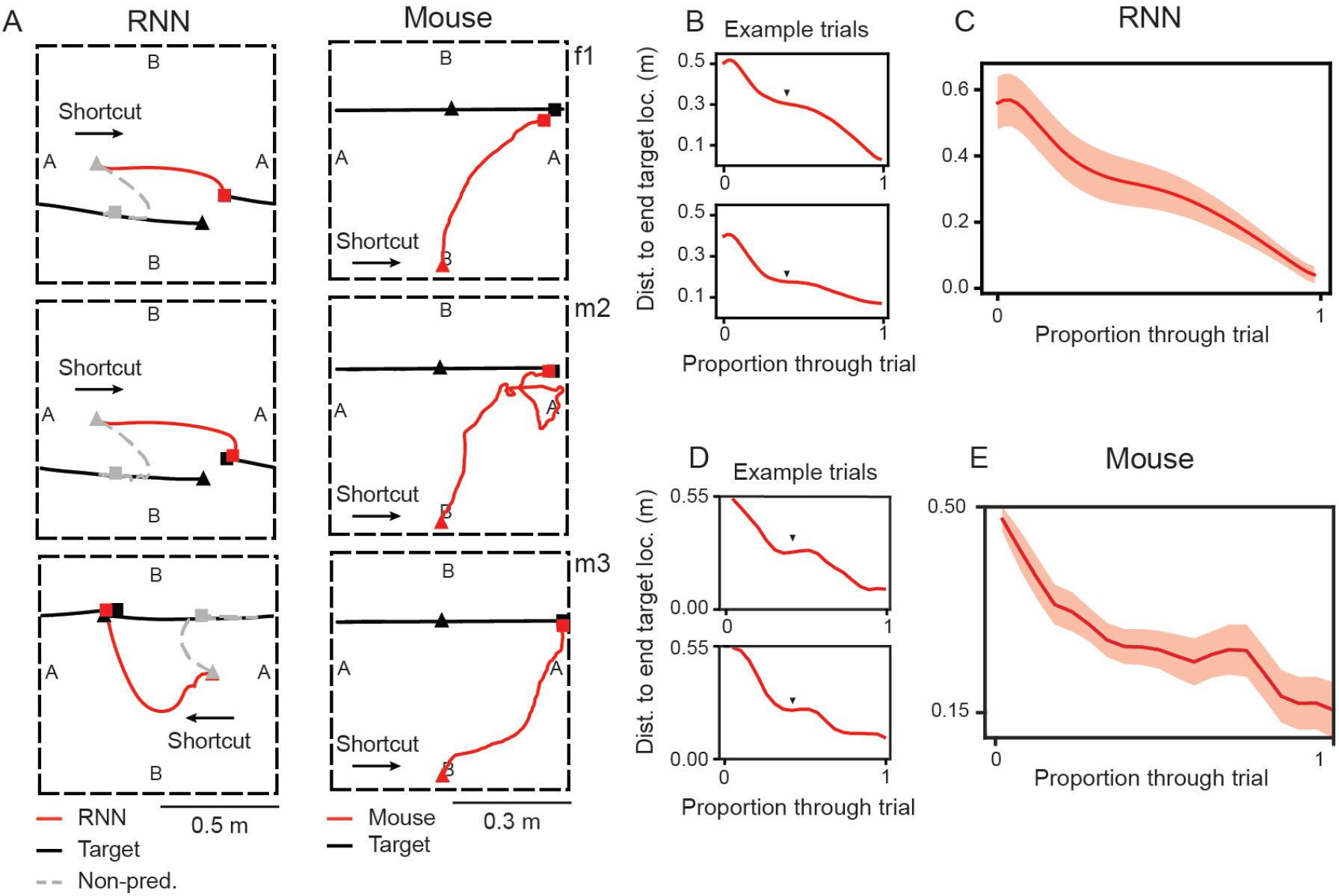
RNNs and mice learn to predictively pursue target in environment with periodic boundaries. (A) Examples of RNN and mouse trajectories for CTs in an environment with periodic boundary conditions (dashed black lines). “A” and “B” denote sides of the environment that are connected. Each row in the mouse column is a distinct animal (f1, m2, m3). (B) Distance of the RNN-agent to the end location of the target-agent, as a function of trial time, for example CT trials. Black triangles denotes period of “waiting”, where distance does not change. (C) Average distance of the RNN-agent to the end location of the target-agent, as a function of trial time, across all CTs starting in the same location. (D) Distance of mouse (m2) to the end location of the laser pointer target, as a function of trial time, for example CT trials. Black triangles denotes period of waiting. (E) Average distance of mouse (m2) to the end location of the laser pointer target, as a function of trial time, across CTs.

From these observations, the RNN models make specific and testable predictions of how an agent (even a biological one) might conduct predictive pursuit on CT trials in an environment with periodic boundaries. In particular, the pursuer should move opposite to the target-agent and then wait, in anticipation of its emergence from the opposite boundary. We test these two predictions by extending the experimental paradigm developed in Alexander et al. (2022) [7] and training mice to perform pursuit tasks in such an environment (see Appendix F for details). As predicted by the RNN, we find that mice adopt the same anticipatory behavior on CT trials as the RNN, wherein they run towards the boundary where the target-agent will appear (Fig. 5A, right column) and wait to capture it when it appears (Fig. 5D; black triangles, also see Fig. 5E for trial averages and Fig. S3 for all mice data). These results provide evidence of similar behavioral strategies emerging from task and environmental constraints across both animal and RNN models of pursuit, highlighting the potential of RNNs to generate experimentally testable predictions of behavior in novel pursuit settings.

### Predictive pursuit emerges in high-dimensional RNNs

Given that pursuit can be performed either predictively (Fig. 3) or reactively (Fig. S2), we next seek to understand what aspects of the RNN model architecture decide the nature of the learned strategy. We hypothesize that the degree of predictive behavior scales with computational resources, such that stronger predictive behavior emerges in networks with higher-dimensional neural codes. To test this hypothesis, we turn to RNNs with low-rank weight matrices [29, 30]. These networks have the same number of recurrent units as the original (“full-rank”) model, but have weights that are increasingly interdependent as the rank decreases, reducing the number of free parameters (see Appendix A.2 for details). In low-rank RNNs, the intrinsic dimensionality of the neural code is bounded above by the rank of the weight matrix [33], therefore manipulating the rank of the network provides a method for controlling the number of coding (“latent”) variables without changing the network size.

We find that RNN models with varying ranks (ranging from 10 to 1000) can learn to perform pursuit well, on both RTs and CTs, although higher-rank models achieve slightly better pursuit accuracies (Fig. 6A). Increasing the rank of the RNNs leads to an increase in the effective dimensionality of the network activations and latent variables, as measured by the number of principal components needed to explain 95% of the variance (Fig. 6B). This is confirmed when using participation ratio as an alternative dimensionality estimate (Fig. S4). These observations are in direct contrast to previously studied low-dimensional tasks, in which latent dimensionality remains constant with increasing ranks [33], or the notion that the dimensionality of computation is set by the complexity of the task [53]. Instead, this increased dimensionality suggests that the RNN actively exploits additional computational resources as rank increases.

**Figure 6:**
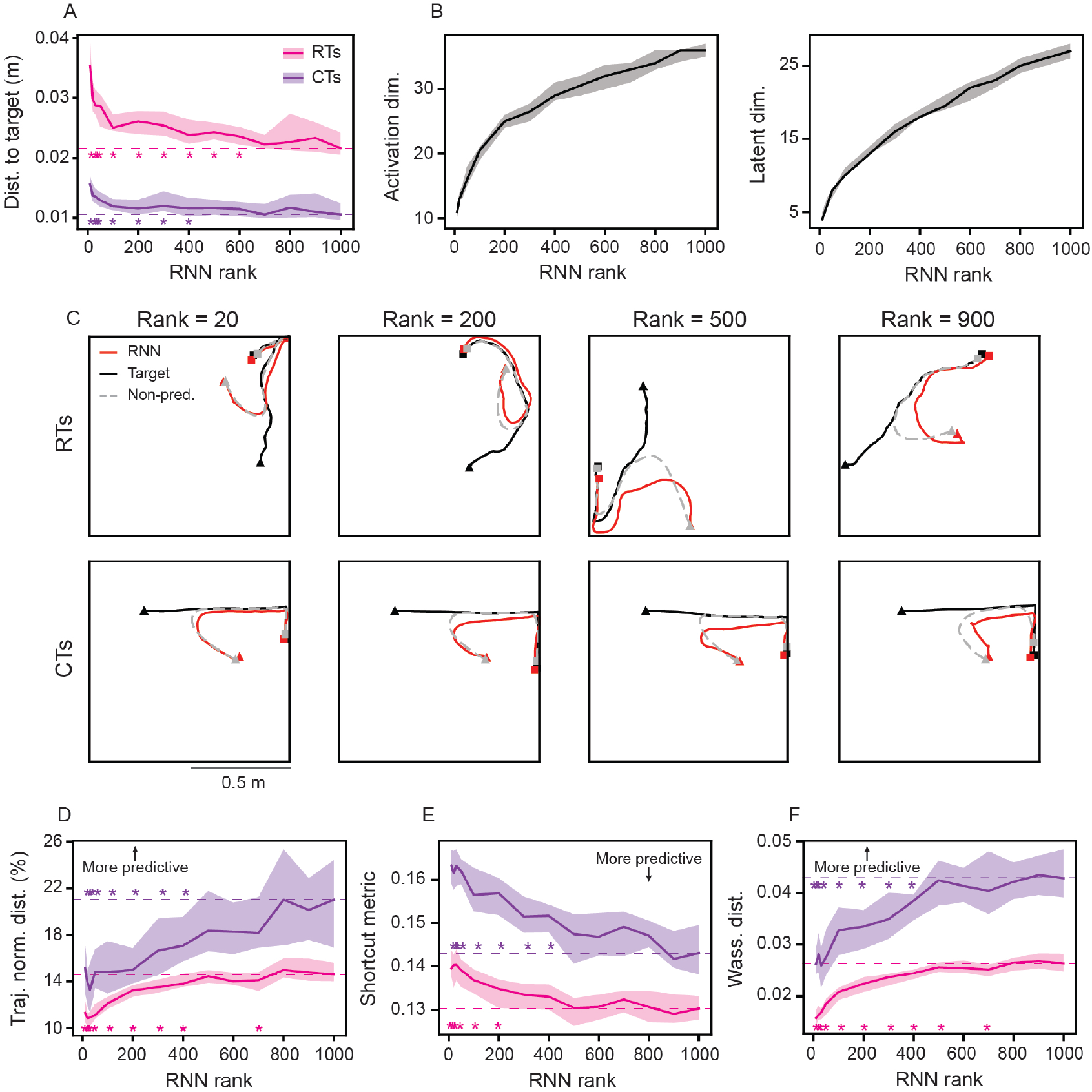
Increasing RNN rank leads to greater predictive behavior. (A) Distance to end location of target-agent for RNNs trained with varying ranks. Solid line is mean and *±* shaded area are standard deviation across 30 independently trained networks. Stars denote ranks where two-sample Kolmogorov-Smirnov test showed significance with respect to rank 1000 performance (*p* < 0.05; dashed lines denote rank 1000 model median value). (B) Dimensionality of unit activations (left) and latent variables (right), computed as the number of PCA dimensions needed to explain 95% of the variance, for RNNs trained with varying ranks. Solid line is mean and shaded area are *±* standard deviation across 30 independently trained networks. (C) Example RT and CT trials for RNNs with different ranks. (D) Normalized distance between RNN and non-predictive control model trajectories, as a function of RNN rank. (E) Shortcut metric, as a function of RNN rank. (F) Wasserstein distance between RNN and non-predictive control model shortcut metric distributions, as a function of RNN rank. (D)–(F) Solid line is mean and shaded area is *±* standard deviation across 30 independently trained networks. Stars denote ranks where two-sample Kolmogorov-Smirnov test showed significance with respect to rank 1000 model distribution (*p* < 0.05; dashed lines denote rank 1000 model median values).

To understand the effects of a higher dimensional code on the RNN behavior, we next inspect individual RT and CT trials. We find that increased rank leads to larger deviations between RNN and non-predictive control model trajectories (Fig. 6C). For instance, when the RNN model has rank 20, the RNN-agent heads towards the location of the target-agent during a representative CT trial, generating a trajectory that is almost identical to the non-predictive control agent (Fig. 6C, left column; compare red and gray lines). In contrast, when the RNN model has rank 900, it generates a trajectory that preemptively heads towards the wall that the target-agent is moving towards, leading to a larger deviation from the non-predictive model (Fig. 6C, right column; compare red and gray lines). These differences in predictive behavior of the RNNs, as a function of rank, generalize across both RT and CT trials (Fig. 6D-F). In particular, increasing RNN rank leads to greater normalized distance between RNN and non-predictive control model trajectories (Fig. 6D). These deviations reflect the emergence of predictive strategies, as shortcut metrics of RNNs decrease with increased ranks (Fig. 6E) and their distributions become increasingly distant from those of the non-predictive control models (Fig. 6F). This rank-dependence is absent when RNNs are trained to minimize the average distance to the target rather than the final distance, in which case all networks, irrespective of rank, learn more reactive strategies (Fig. S5). In addition, for a fixed rank, we find similar predictive behaviors in networks when varying the number of units (Fig. S6), implying that the dimensionality of the latent variables, not the number of units, is associated with the predictive behavior. Overall, these results demonstrate that predictive pursuit emerges in high-dimensional networks. Importantly, this is not a generic property of high-rank RNNs, but arises specifically when a predictive strategy is useful.

### High-dimensional networks learn to represent allocentric information

Having shown that the degree of predictive behavior scales with the dimensionality of the neural code, we next set out to extract the qualitative changes in neural representations in high-rank networks that may underlie this emergent strategy. Specifically, we examine how well distinct task features can be linearly decoded from the population activations of RNN models trained with varying ranks (see Appendix G for more details). This linear decoding acts as a proxy for how well these features are represented by the RNN models and can be used to extend our retrospective and prospective coding analysis in Fig. 2 from individual units to the entire RNN population.

Given that removing ETUs led to significantly worse pursuit performance (Fig. 2), we first decode current, retrospective, and prospective egocentric target distances from population activations. Intuitively, even an agent performing a purely reactive pursuit would need to keep track of the distance to the target-agent. In-line with this intuition, the distance can be decoded with high accuracy across RNN models of all ranks (Fig. 7A). While the lowest decoding error was achieved for the current egocentric target distances, the decoding was highly accurate for both past and future values. The qualitative uniformity of the decoding performance suggests that the use of egocentric target coding as a pursuit strategy is common across all RNN model ranks, and therefore is unlikely to be the driver of the emergence of predictive behavior that we observe in higher-rank models.

**Figure 7:**
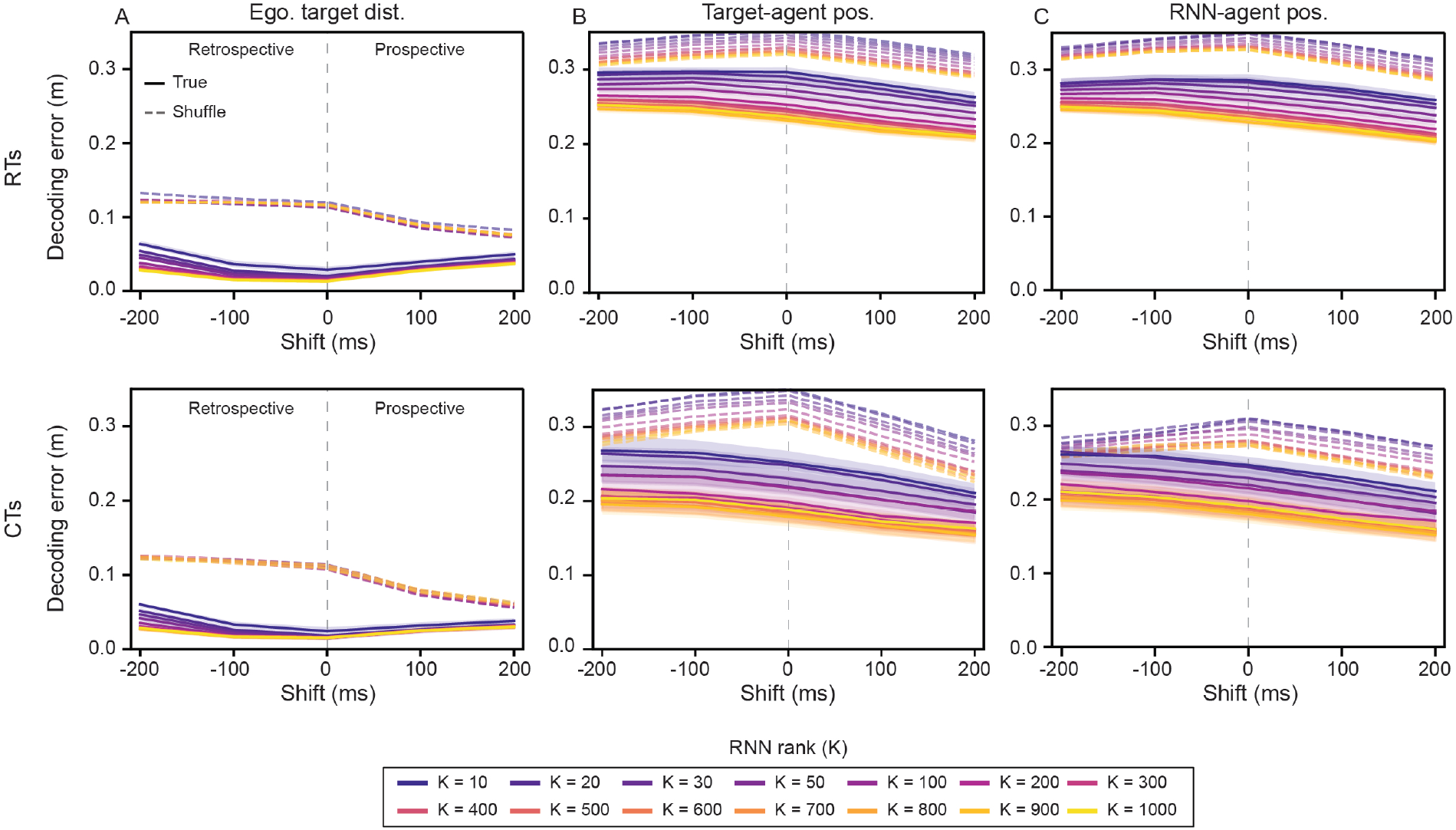
Decoding accuracy of RNN-agent and target-agent position increases with increasing RNN rank. (A) Decoding error on RTs (top row) and CTs (bottom row) of egocentric target distance, for trained RNNs of varying rank, across prospective and retrospective shifts. Gray dashed line denotes no shift (current value). (B)–(C) Decoding error on RTs and CTs of target-agent and RNN-agent position, for trained RNNs of varying rank, across prospective and retrospective shifts. Solid lines denote median and shaded area denotes *±* standard deviation, across 30 independently trained RNNs. Dashed lines denote median shuffled distribution.

Next, we turn to allocentric features (i.e., positions of the target-agent and the RNN-agent). Notably, the RNNs are not explicitly given position information in allocentric coordinates (except the initial positions). Therefore, any allocentric position estimate that exists in the RNNs must be constructed by integrating velocity inputs over time. Unlike egocentric target distance, allocentric variables exhibit a strong rank dependence, with increasing rank leading to better accuracies (Fig. 7B, C). Furthermore, positions of both agents can be decoded with higher accuracies in CTs than in RTs (Fig. 7B, C; compare top row with bottom row), consistent with the repeated structure of CTs, which constrains trajectories and thereby makes positions more predictable. In addition, for both agent’s positions, decoding error decreases with prospective shift and increases with retrospective shift (Fig. 7B, C), indicating that population activations preferentially reflect predictions rather than past states. We observe similar results when we quantify the decoding performances via explained variances (Fig. S7).

We find that the metrics quantifying shortcut-like pursuit are strongly associated with the decoding error of target-agent and RNN-agent positions (Fig. S8; top row and middle row – absolute Pearson’s correlation coefficients 0.64 ≤ |*r*| ≤ 0.81). The correlation between predictive behavior and decoding error of egocentric target distance is also statistically significant, though slightly weaker (Fig. S8; bottom row – 0.37 ≤ |*r*| ≤ 0.60). For the RNNs trained to minimize the average distance between the target-agent and the RNN-agent across the entire trial, we find only weak correlations between decoding error of all three features and predictive behavior (Fig. S9; 0.06 ≤|*r*| ≤ 0.41).

The association between rank and predictive pursuit, combined with these decoding results, suggests a mechanism through which higher-dimensional codes collectively drive the shift toward predictive strategies. The emergence of predictive pursuit in high-rank RNNs is thus accompanied by an increased capacity to encode allocentric information, providing a key experimentally testable signature for identifying brain regions that support predictions during active pursuit.

## Discussion

### Development of an RNN framework for examining predictive chasing behaviors

Predictive pursuit of a dynamic target is an evolutionarily important and widespread behavior [2–9]. Understanding the neural computations that support this behavior offers the potential to gain insight on fundamental cognitive processes, such as planning and forming internal models [1]. Studying these ethologically relevant computations in a controllable laboratory setting is challenging, and prior experimental paradigms have either prioritized ethological relevance over control of the target-agent [54, 55] or prioritized control of the target-agent over ethological relevance [8, 9]. The recent development of tasks that involve unrestrained rodents pursuing (or avoiding) target-agents controlled by experimenters has demonstrated their ability to develop complex, long-time scale strategies that involve predictive decisions [7, 40, 56]. Simultaneous electrophysiological recordings have also provided insight into neural representations that may underlie this behavior (e.g., egocentric target cells) [7, 40].

Motivated by these experimental results, as well as the desire to probe – at a level not yet possible experimentally – the computations that support predictive pursuit, we developed an RNN model and trained it to chase a moving target. We find that, when using a loss function that penalizes the distance between the RNN-agent and the target-agent at the end of the trial, the RNN model generates trajectories in a predictive manner, which is increased when the target-agent has trajectories with repeated structure (Fig. 3). Leveraging knowledge of the target-agent’s behavior is experience dependent, with increased exposure leading to increased anticipatory behavior (Fig. S1). This is in contrast to RNN models trained to minimize the average distance between the RNN-agent and the target-agent (Fig. S2), highlighting how the intrinsic goal (i.e., loss function) shapes the learned behavioral strategy.

### Emergent egocentric target representations in RNNs trained to pursue moving targets

Previous experimental works have identified egocentric target cells in PPC and RSC during pursuit task [7, 40]. We find that the RNNs develop units with analogous tuning (Fig. 2), despite receiving velocity inputs anchored to the allocentric coordinate system of the simulated arena. This demonstrates that the RNNs have learned to perform an allocentric-to-egocentric transformation, a process which has been proposed to be critical for spatial navigation tasks [57–60] and has been found to support multi-agent path integration in RNNs [46]. Accordingly, future work could use RNNs to explore population level computations underlying coordinate transformations across spatial reference frames.

While experimental recordings have identified ETCs in visual association cortices, they have not yet shown that they play an important, *causal* role in pursuit (though see [61]). As is the case with many functionally-defined representations of this nature, it is in fact unlikely that a total ablation of solely egocentric target sensitive neurons is even possible with current experimental techniques. Thus, our RNN framework provides a unique opportunity to establish a theoretical causality between these representations and pursuit behaviors of simulated agents. By ablating the units with the highest egocentric target information, we find a significantly greater degradation of performance than randomly removing units (Fig. 2C). Thus, in our RNN model, information about the target, in the transformed egocentric reference frame, is useful to pursuit performance.

### RNNs develop internal models that subserve predictive pursuit

To explore the capability of the RNNs to develop and maintain an internal representation of the target-agent’s position, we trained on trajectories where a percentage of the inputs to the RNN model were “masked.” This loosely mimics the presence of obstacles that occlude the target-agent, as well as changes in attention. Even when the inputs are masked 50% of the time, the RNN-agent pursues the target-agent along shortcut-like trajectories (Fig. 4), demonstrating its ability to leverage an internally maintained prediction of the target-agent’s position. These results motivated us to consider whether the RNN model could similarly learn to perform predictive pursuit in a more complex environment. To this end, we trained the RNNs to pursue the target-agent in an environment with periodic boundaries, introducing a non-Euclidean geometry to positions in the environment. We find that the RNNs learn to anticipate the target-agent’s trajectory from one boundary to another on CTs, with the RNN-agent waiting near its starting location, until the target-agent has emerged (Fig. 5A–C).

After observing the strategy developed by the RNN, we tested whether similar behaviors emerged in biological agents by examining target pursuit in mice under conditions that also featured periodic boundaries. To this end, we developed a behavioral-closed-loop target presentation system that allowed precise control of target movement, teleportation, and interception. We demonstrate that, like the RNN, mice ignore target movement towards the “disappearing” boundary and instead navigate directly to the “reappearing” edge, where they wait to efficiently intercept (Fig. 5A, D, E). The similarity in capture strategies highlights rich behavioral-level agreement between real and artificial agents. In addition, this suggests that our RNN model may be useful for piloting possible experimental paradigms that can later be tested in the lab.

### Low-rank RNNs are sufficient for successful pursuit, but higher dimensional computations are required to mimic predictive behaviors observed in biology

Having shown that the RNN model learns to perform an allocentric-to-egocentric transformation and develops an internal representation of the target-agent’s position, we asked whether and how the computational capacity of the RNNs enable these complex processes to emerge. To probe this, we trained RNN models of varying rank and examined the learned behavioral strategies. We find that predictive trajectories are learned in high-rank RNN models, with lower-rank RNN models generating trajectories that are significantly more similar to the non-predictive control model (Fig. 6). We further find that even though RNNs are capable of conducting pursuit without developing allocentric information, allocentric representations of the RNN-agent and the target-agent emerge in networks with high-dimensional neural codes (Fig. 7), reminiscent of “social” place coding [62, 63] that has been found to emerge in pursuit tasks [64]. That this happens, despite RNN models of low-rank having similar performance on the pursuit task (especially on CTs), is indicative of the development of the complex computational properties that underlie predictive pursuit in high-rank RNNs. This is, to our knowledge, the first time high-dimensional network structure has been shown to be necessary to generate a given low-dimensional behavioral regime (the network output is only two-dimensional). In addition, it highlights the fact that, in certain scenarios, predictive behavior emerges not solely because it is beneficial to achieve performance on a task [65], but because the network structure has sufficiently large capacity to support its development.

Collectively, our results demonstrate that predictive pursuit is a rich cognitive process and that use of RNN models enable the testing of novel experimental paradigms (i.e., pursuit in a periodic boundary conditions environment), the causal probing of the computational role of putative functional classes (i.e., the ablating of ETUs), and the identification of underlying computational principles (i.e., the necessity of high-dimensional network structure).

### Limitations

Our RNN model assumes true allocentric movement direction and speed of the target-agent as inputs, and true allocentric movement direction and speed of the RNN-agent as outputs. Incorporating noise in the estimate of self-motion [66] and target-agent motion, as well as including other types of inputs (e.g., visual-scene information) could bring our RNN model closer to biological systems. However, we note that the RNNs being able to continue to perform predictive pursuit when inputs were masked (Fig. 4) provides evidence that it would be able to perform well with noisy movement estimates. The steering of the RNN-agent is controlled by only two outputs, the RNN-agent’s movement direction and speed. This greatly simplifies the transformation that occurs between associations areas (e.g., PPC, RSC) and primary motor areas. In addition, it makes the RNN model sensitive, as any perturbations to weights or activations can directly cause changes to the RNN-agent’s behavior. While this could cause the low-rank RNNs to be less maneuverable, we note that the high performance on CTs and RTs, across ranks (Fig. 6A), suggests that this is not the major driver of the behavioral differences we see between low-rank and high-rank networks. Adding redundancy in the output layer (e.g., multiple outputs that are averaged together to determine RNN-agent movement direction and speed) could enable more robust steering. The supervised training of our RNN model simplifies aspects that shape rodent behavior, such as motivation and reward. Reinforcement learning (RL), which has been used to study aspects of pursuit-like behavior [65, 67–71], offers a useful framework for capturing these components. While our results demonstrate that supervised learning is sufficient to generate similarities between the RNN model and rodent experiments at the behavior (Fig. 5) and computational level (Fig. 2), training the RNN model using RL offers the ability to generate a broader range of behavioral and computational solutions.

### Future directions

Our work opens the door for a number of exciting new directions to computationally study predictive pursuit. First, the target-agent we train our RNN to pursue is not “aware” it is being chased and therefore, does not perform any evasive strategies. While this may be appropriate for modeling some kinds of predatory-prey interactions (e.g., bats hunting mosquitoes [4–6]), many interesting scenarios involve both the predator and the prey exhibiting adaptive and predictive behavior. Thus, understanding how the RNN model can learn to pursue an “intelligent” target-agent and how the anticipatory behavior of the target-agent shapes the needed level of prediction by the RNN will provide a deeper understanding of the neural computations that may support predictive pursuit. Second, the environment the RNN-agent and target-agent are constrained to is empty and has a simple geometry and topology. Studying the representations that emerge in RNNs trained to perform pursuit in cluttered environments [56, 65] and environments with complex topology (e.g., tree branch-like graphs that jumping spiders hunt on [72]) will enable an understanding of how they are shaped by properties of the environment and its affordances. And third, we have focused on predictive pursuit, but many organisms must be able to perform predictive evasion [56, 73, 74]. The RNNs can be made to avoid the target-agent by flipping the sign of the loss function, however whether and how they can learn to do this in a predictive manner is not known. Comparing the representations that emerge in RNNs trained to perform pursuit with the representations that emerge in RNNs trained to perform evasion can provide insight into what computations are generally useful for prediction in navigation based environments, and which computations are useful for specific roles (e.g., predator, prey).

## Acknowledgments

This research was supported in part by grant NSF PHY-2309135 and the Gordon and Betty Moore Foundation Grant No. 2919.02 to the Kavli Institute for Theoretical Physics (KITP). A.S.A, and X.L., and M.G.C. were supported by NIH R00 NS119665. A.S.A. and M.G.C. were supported by the U.S. Department of Energy, Office of Science, Advanced Scientific Computing Research (ASCR) program as part of the DOE Collaborative Research in Computational Neuroscience (CRCNS) Program. Author contributions: W.T.R., F.D., and A.S.A. designed the study. W.T.R., F.D., and X.L. conducted all experiments. W.T.R., F.D., M.G.C., X.L., and A.S.A. analyzed the data. W.T.R., F.D., and A.S.A. wrote the paper. All authors assisted with revision of the manuscript. Competing interests: The authors declare that they have no competing interests. Data and materials availability: All data needed to evaluate the conclusions in the paper are present in the paper and/or the Supplementary Materials. Additional data related to this paper may be requested from the authors.

## A Task and the RNN models

### A.1 RNNs as pursuit agents

We consider the setting where two agents (“RNN-agent” and “target-agent”) traverse a square environment through finite length paths. The target-agent starts at position **z**_target_(0) = (*x*_target_(0)), *y*_target_(0)) ^⊺^ ∈ [−*L/*2, *L/*2] *×* [− *L/*2, *L/*2] and subsequently moves to positions **z**_target_(0) → **z**_target_(1) → … → **z**_target_(*T* ), where the path is sampled using a model of motion that is biased towards straight paths and avoidance of walls [19, 27] (Fig. 1A). This path can alternatively be defined by **z**_target_(0) and a sequence of movement directions, *θ*_target_(*t*) ∈ [−*π, π*], and speeds, *v*_target_(*t*) ∈ [0, ∞), such that

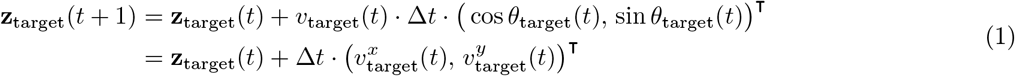

for small Δ*t* ∈ ℝ^+^ and *t* = 1, …, *T*, and where 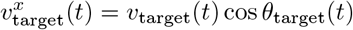 and 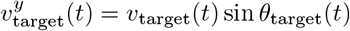.

The RNN-agent similarly starts at a random position in the environment, **z**_RNN_(0) = (*x*_RNN_(0), *y*_RNN_(0) ^⊺^ ∈ [− *L*/2, *L*/2] × [−*L*/2, *L*/2] (Fig. 1A). We train the RNN to generate a sequence of movement directions, *θ*_RNN_(*t*) ∈ [−*π, π*], and speeds, *v*_RNN_(*t*) ∈ [0, *v*_max_), such that the trajectory of the RNN-agent, **z**_RNN_(0) → **z**_RNN_(1) → … → **z**_RNN_(*T* ), is as close to the target’s end position as possible (Fig. 1C). This is enforced by the loss function,

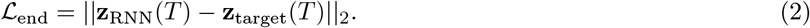

To learn an RNN model that is able to perform pursuit, we utilize the following architecture (Fig. 1B). Inputs, in the form of 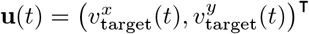, are projected into the recurrent layer of an RNN via weights **W**^in^ ∈ ℝ^*N ×*2^, where *N* is the number of hidden units. The recurrent units process the input via the weights **W**^rec^ ∈ ℝ^*N ×N*^, and drive activations in output units via the weights **W**^out^ ∈ ℝ^2*×N*^ . More explicitly, the dynamics of the RNN model are given by

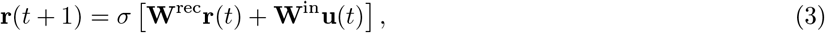

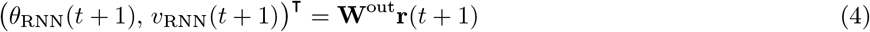

where **r**(*t*) ∈ ℝ^*N*^ are the activations of the recurrent units at time *t, σ*(·) is the activation function, and *θ*_RNN_(*t*) and *v*_RNN_(*t* + 1) are the generated movement direction and speed for the RNN-agent at time *t*. To prevent the RNN from using unreasonably large instantaneous speeds to perform pursuit, we threshold the outputted *v*_RNN_(*t* + 1), setting *v*_RNN_(*t* + 1) = min{*v*_RNN_(*t* + 1), *v*_max_)}, where *v*_max_ ∈ ℝ^+^. Given the output *θ*_RNN_(*t*), *v*_RNN_(*t*), we can generate the trajectory of the RNN-agent, **z**_RNN_(0) → **z**_RNN_(1) → … → **z**_RNN_(*T* ) by using *θ*_RNN_(*t* + 1) and *v*_RNN_(*t* + 1) to get 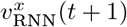 and 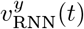. Training can then be achieved by using Eq. 2 as the objective function.

In addition to the movement direction and speed of the target agent, in order to successfully achieve pursuit, it is necessary to know where the RNN-agent and target-agent are at the beginning of a trial. Therefore, we additionally learn an embedding of the **z**_target_(0) and **z**_RNN_(0) to the initial state of the RNN, via the weights **W**^back^ ∈ ℝ^*N ×*4^. That is, **r**(0) = **W z**_RNN_(0), **z**_target_(0) . All parameters associated with the RNN architecture and all hyper-parameters associated with training the RNN are presented in Table S1.

### A.2 Low-rank RNNs

To constrain recurrent connectivity, we replace the full recurrent weight matrix with a rank-*K* factorization. Specifically, instead of learning **W**^rec^ ∈ ℝ^*N ×N*^ freely, we parameterize it as

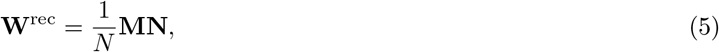

where **M** ∈ ℝ^*N ×K*^, **N** ∈ ℝ^*K×N*^, and *K* ≤ *N* is the prescribed rank. By construction, rank(**W**^rec^) ≤ *K*. The factor 1*/N* ensures stable scaling of recurrent contributions as *N* varies. Under this parameterization, the recurrent update becomes

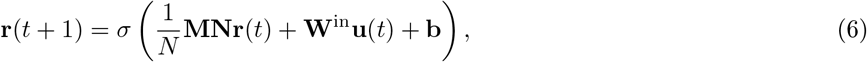

with the same input, output, and loss definitions as in the full-rank case. The recurrent contribution depends only on the projection

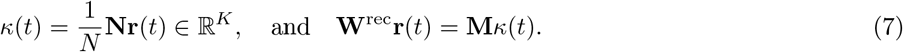

**Table S1:** Parameters used to train RNNs on pursuit. Unless otherwise noted, the parameters presented below were used to train the RNN on the pursuit task.

| Parameter | Value |
| --- | --- |
| Epochs | 100 |
| Batch size | 200 |
| Batches per epoch | 1000 |
| Path length ($T$) | 50 |
| $\Delta t$ | 0.02 |
| Arena length ($L$) | 1.0 m |
| Learning rate | $2.5^{-5}$ |
| $N$ | 1024 |
| Activation | ReLU |
| Weight decay | $10^{-4}$ |
| Optimizer | Adam |
| Percent CTs | 25% |
| $v_{\max}$ | 0.0225 |

Thus, the recurrent drive lies in the *K*-dimensional column space of **M**, constraining network dynamics to evolve within a low-dimensional subspace when *K* ≪ *N*. The number of recurrent parameters is reduced from *N* ^2^ to 2*NK*.

Parameters were initialized independently as Gaussian random variables:

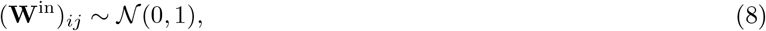

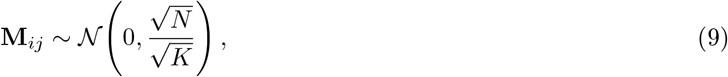

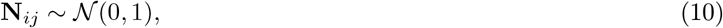

so that the recurrent weights are effectively initialized 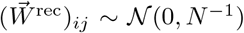. For *N, K* → ∞, this initialization corresponds to the edge of chaos [38, 75], which has been known as a fruitful initialization in the broader deep learning literature [76]. We trained networks across a range of ranks *K* ∈ {10, 20, 30, 50, 100, 200, …, 1000}, with ten independent random initializations per rank and no weight decay. We used a learning rate of 10^−4^. We trained for 100, 000 epochs, using a single batch per epoch, matching the total training time as in Table S1 but keeping a more frequent record of the loss function values. All other training hyperparameters were held fixed.

## B Characteristic trajectories

Inspired by the experimental paradigm developed in Alexander et al. (2022) [7], we train RNNs on trajectories that are sampled from two different families. The first, which are referred to as pseudo-random trials (RTs), are generated using the same approach as has been described elsewhere [19, 27]. The second, which are referred to as characteristic trials (CTs), are generated using the following procedure. First, the target is randomly assigned to one of four starting locations: 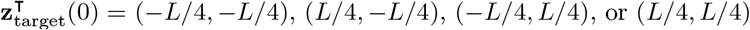, where *L* is the length of the square environment and the environment is defined by [− *L/*2, *L/*2] × [− *L/*2, *L/*2]. Once the starting location of the target-agent is chosen, then the location of the RNN-agent and the starting movement direction of the target-agent are determined. In particular,

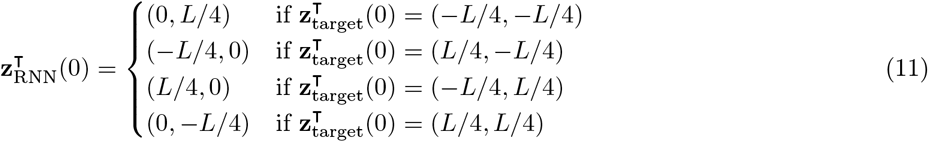

and

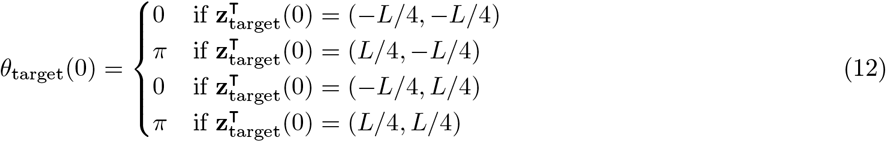

The target then moves along trajectories that are updated by sampling speeds from the same distribution as the RTs and sampling changes to the head direction from a normal distribution, where the standard deviation in the distribution for CTs is 1 radians/second, as compared to 11.52 radians/second for RTs. Finally, if the target-agent gets sufficiently close to the wall, the movement direction changes by +*π/*2 radians if the target-agent is in the upper-right or lower left-hand quadrants, and the movement direction changes by − *π/*2 if the target-agent is in the upper-left or bottom-right quadrants. The resulting affect of this turn is to make the target-agent move along the wall, towards *x* = 0. This creates an “L” shaped-like trajectory (Fig. 1F, black lines).

## C Egocentric target units

Electrophysiological recordings of rodents performing pursuit have found cells in RSC and PPC that encode the position of the target-agent in the egocentric coordinate system [7, 40]. We asked whether the RNNs developed similar tuning. To investigate this, we followed similar steps to previous work on analyzing egocentric target and border coding [7, 39]. First, we sample 10000 RT trials, computing the distance and angle between the RNN-agent and the target-agent at each time-point. We define this angle as

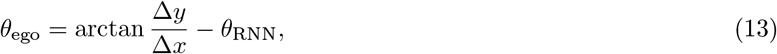

where *θ*_RNN_ is the movement direction of the RNN-agent, Δ*x* = *x*_target_ − *x*_RNN_, and Δ*y* = *y*_target_ − *y*_RNN_. Care needs to be used when taking the arctan to ensure the sign of the output is correct.

We similarly save the activations of all units in the RNN at each time-point. We then split the 10000 RT trials into two subsets of 5000 distinct trials and construct the egocentric target ratemaps by binning distance and angle, and averaging the activity associated with time-points that fall within each bin. Because the RNN was trained to pursue the target-agent, there were few time-points with large distance. Following previous analysis of neural data, we set a maximum distance, *d*_max_ = 0.4 m. Any time-point with distance greater than *d*_max_ was discarded and not used for analysis. We denote the two constructed egocentric target ratemaps of unit *i* by 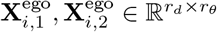, where *r*_*d*_ is the number of bins for distance and *r*_*θ*_ is the number of bins used for angle. We then compute the mean resultant length (MRL) of these egocentric ratemaps [39]. The MRL is given by

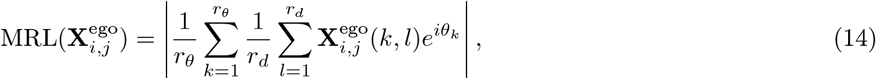

where *θ*_*k*_ is the value corresponding to the center of the *k*^th^ angular bin (ranging from − *π* to *π*).

For each unit in the RNN, we have two MRL values, 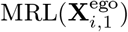 and 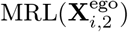, corresponding to the two separate sets of trials. We assess the robustness of the egocentric coding by computing

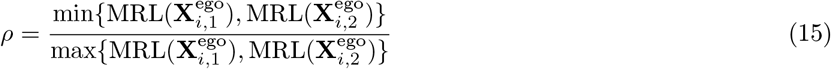

A unit is considered robust is this ratio is greater than 0.95. For all units that are robust, we construct the full egocentric target ratemap, 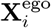, where we use all 10000 trials. The corresponding MRL is 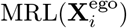.

To assess which robust units have significant egocentric target tuning, we generate 100 shuffled egocentric target ratemaps 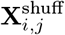 by permuting the unit activations, with respect to the distances and angles to the target-agent, on each trial. We then consider the distribution of MRL values computed on all shuffled egocentric target ratemaps. We identify units with significant egocentric target tuning as those with 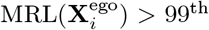 percentile of the shuffled distribution (Fig. 2B). Units that are both robust and have significant egocentric target tuning are classified as egocentric target units (ETUs).

### C.1 Ablating egocentric target units

Having found units that are classified as ETUs (Fig. 2A, top row), we asked whether they were used by the RNN to perform pursuit. To test this, we perform targeted ablations [27, 28]. This was achieved ordering ETUs by their MRL and ablating the top *pN* units, where *p* ∈ [0, 1] and *N* is the total number of units in the RNN, by setting 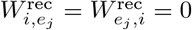, where *e*_*j*_ is the *j*^th^ ETU and 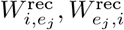 are the in-going and out-going recurrent weights to unit *e*_*j*_. Setting these weights to 0 effectively removes unit *e*_*j*_ from the RNN.

To identify the significance of removing ETUs (as compared to other units), we perform two additional ablation experiments. First, we randomly select *pN* units to ablate. For each *p*, we perform 25 random selections. This enables us to assess whether removing ETUs leads to a more significant effect on pursuit performance than randomly removing units. Given that approximately 25% of RNN units were found to be ETUs, this is a strong baseline [28], as randomly selected units will be ETUs with non-trivial probability. Therefore, we also examine how ablating the RNN units with the lowest MRL affects pursuit performance. In this case, we sort all units by their MRL (not just ETUs) and remove the bottom *pN* units (Fig. 2A, bottom row). This allows us to examine how important egocentric target coding is to pursuit performance.

## D Non-predictive control model

To understand the behavior of the RNN model, we compare its performance to a control model that, by definition, is non-predictive. To do this, we consider the locally optimal solution of moving, at each time step, in the direction of where the target-agent is. More precisely, if **z**_target_(*t*) is the current position of the target-agent and **z**_non-pred_(*t*) is the current position of the non-predictive-agent, then

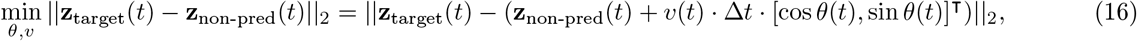

is achieved by

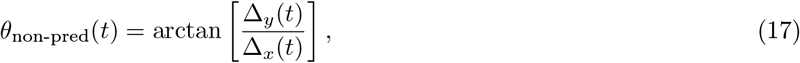

where [Δ_*x*_(*t*), Δ_*y*_(*t*)]^⊺^ = **z**_target_(*t*) −**z**_non-pred_(*t*). Care has to be taken to add *π* or 2*π* to Eq. 17, depending on the signs of Δ_*x*_ and Δ_*y*_.

With these movement directions, the non-predictive control model generates a trajectory via

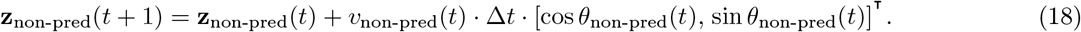

Importantly, if *v*_non-pred_(*t*) is unconstrained, then **z**_non-pred_ can catch-up instantly to **z**_target_. Thus, to ensure a fair comparison, we take *v*_non-pred_(*t*) to be the same as the velocity of the RNN. That is, *v*_non-pred_(*t*) = *v*_RNN_(*t*). This allows us to identify how the RNN differs from the non-predictive control model in terms of its planning ability through trajectory direction it takes.

Note that this is similar to the baseline with which Yoo et al. (2020) [8] compared their non-human primate results with. However, this is a simpler model that captures a fully non-predictive strategy.

### D.1 Shortcut metric

To quantify the ability of the RNN to perform anticipatory behavior, we developed the following “shortcut metric”. To this end, we measure the average distance between the RNN’s trajectory and the trajectory with the minimum path length (i.e., the “globally optimal” solution). The globally optimal trajectory is given by the linear interpolation between the starting position of the RNN-agent, **z**_RNN_(0), and the end position of the target-agent, **z**_target_(*T* ),

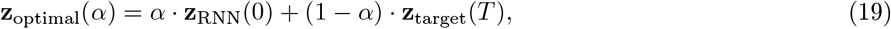

where *α* ∈ [0, 1] describes the interpolation.

Let *ℓ*_RNN_(*t*) be the minimum distance between the RNN-agent’s trajectory at time *t* and the globally optimal trajectory. That is,

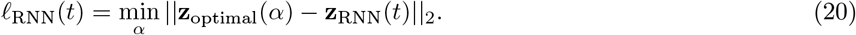

We define our shortcut metric, *ε*_RNN_, to be the mean distance between the RNN-agent’s trajectory and the globally optimal trajectory,

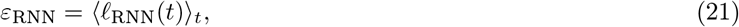

where ⟨·⟩_*t*_ is the mean taken over all time steps in the trajectory.

We analogously define *ε*_non-pred_. Comparing the distribution of *ε*_RNN_ and *ε*_non-pred_ across many sampled trajectories enables us to quantify statistically significant anticipatory behavior.

## E Average distance loss

To understand to what extent the predictive behavioral outputs of the RNN model are due to the loss function penalizing the *end* distance between the RNN-agent and the target-agent, we train RNN models using an alternative loss function that penalizes the *average* distance between the RNN-agent and the target-agent. In particular, where as ℒ_end_ (Eq. 2) was a function only of ||**z**_RNN_(*T* ) − **z**_target_(*T* )||_2_, the average loss function is given by

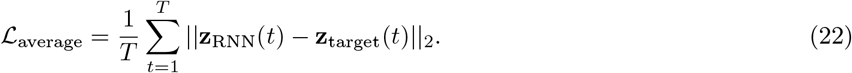

## F Periodic boundary condition environment

To test the capacity of RNN models to learn to perform predictive pursuit, we developed an environment with periodic boundary conditions. That is, we connected the left and right walls, as well as the top and bottom walls. This gave rise to the environment having the topology of a torus, with the distance between two points not equal to the Euclidean distance. The RNN model’s ability to learn anticipatory behavior on CTs in this environment (Fig. 5A, left), led us to develop a similar environment to test mice in. Below, we discuss each of these environments in detail.

### F.1 RNN experiments

To generate the environment with periodic boundaries in simulation, we made the following modifications to our standard environment and process of sampling target-agent trajectories [19, 27]. First, in the standard environment, the target-agent avoids the boundaries by turning away from them as it nears the wall (modeled as an elastic collision). We remove this effect, so that the target-agent in the periodic boundary condition continues towards the wall. And second, on each update of the position of the target-agent, we enforce periodicity of the environment by checking to see if **z**_target_(*t* + 1) ∈ [−*L/*2, *L/*2] *×* [−*L/*2, *L/*2]. If **z**_target_(*t* + 1) is not in this domain, then we perform the following correction

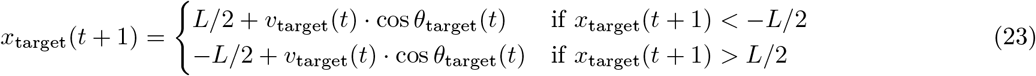

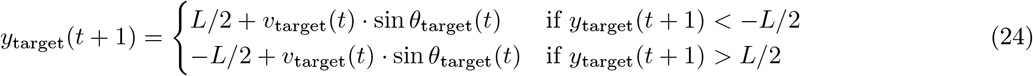

We perform similar corrections to the position of **z**_RNN_(*t* + 1).

Training RNN models to pursue target-agents in this environment, we find low performance on RTs (median distance to end location of target-agent – RNN = 0.34 m). This is presumably because the target-agent is able to quickly “jump” from one part of the environment to the other, and can even make multiple jumps in a short span of time. This makes it challenging for the RNN-agent to catch the target-agent. However, appropriately defined CTs offer the potential for providing enough structure that the RNN model could potentially learn to predictively pursue the target-agent.

Therefore, we sample CTs in this environment, using the same procedure as was done in the standard environment, with the two modifications listed above. Namely, not implementing the boundary avoidance, connecting the left- and right-hand walls, and connecting the top and bottom walls. This leads to CTs that are straight lines (although the presence of noise in the movement direction updates leads to some variability) that wrap around the environment (Fig. 5A, right). We find that the RNN model can learn to perform with high accuracy on CTs (median distance to end location of target-agent – RNN = 0.04 m), demonstrating the ability to learn to pursue in the more complex environment.

### F.2 Rodent experiments

#### Behavioral apparatus and tracking

Rodent pursuit behavior was examined in a square open-field arena (62 × 62 cm) designed to enable the animal to chase a moving visual target. The arena floor consisted of transparent glass, allowing rear-projection of a visual stimulus from a projector positioned beneath the apparatus. The stimulus was a green circular target (diameter, 2.5 cm) displayed on the arena floor. A start box (28 × 19 cm) was attached to one side of the arena and connected to the main arena via a manually operated sliding door. A water spout located inside the start box delivered liquid reward (10 *µ*L) following successful trials via a Bpod behavioral control system running custom MATLAB scripts. To minimize uncontrolled visual landmarks and reduce spatial bias, the arena was surrounded by black-and-white curtains.

Animal behavior was recorded using an overhead camera at 30 Hz. Mouse and target positions were tracked in real time using a trained DeepLabCut model [77], enabling online detection of target interception defined by the proximity between the mouse’s nose and the target center (*<* 1 cm). Upon interception, the target immediately disappeared, and reward delivery was triggered once the mouse returned to the start box. Target presentation, behavioral tracking signals, reward delivery, and event synchronization were coordinated through Bonsai.

#### Target trajectory generation

Target trajectories were generated offline using a custom simulation interface and implemented in two motion regimes: random trajectories (RT) and characteristic trajectories with periodic boundaries (CT-PB). In RT trials, the target followed pseudo-random two-dimensional trajectories with smooth directional transitions while avoiding arena boundaries. Target speed varied between 5 and 30 cm/s and trajectory duration ranged from 7 to 15 s. Each trajectory was generated uniquely across trials to prevent repetition. In CT-PB trials, the target moved at a constant speed (25 cm/s) along a straight trajectory. When the target reached an arena boundary, its position was wrapped to the opposite side while preserving speed and direction, implementing a periodic boundary condition that produced highly stereotyped trajectories across repetitions. CT-PB trials terminated after 1 min or after five full trajectory iterations.

#### Task structure and training

Each trial began with the mouse positioned inside the start box. At trial onset, the target appeared in the arena and the sliding door was opened manually. Target motion along the predefined trajectory was initiated once the mouse’s head entered the arena. A trial was considered successful when the mouse intercepted the target; otherwise, the trial terminated without reward when the trajectory ended. Each session consisted of 30 trials, and animals typically performed one RT session and one to two CT-PB sessions per day. Mice were initially trained on RT trials until stable performance was achieved (≥ 80% interception success), after which CT-PB sessions were introduced to test whether animals could learn and exploit the repeating trajectory structure. Throughout training, animals were maintained under controlled water access (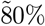 baseline body weight) in accordance with institutional animal care guidelines.

## G Decoding Analysis

To perform our decoding analyses, we focus on fully trained low-rank RNNs on the two types of pursuit tasks, i.e., when the loss function focuses on the final distance (Figs. 7, S7, and S8) and on the trial-averaged distance (Fig. S9). For a given RNN and trial type (random vs characteristic), we generated independent trials and collected the RNN-agent (**z**_RNN_(*t*)) and target-agent (**z**_target_(*t*)) trajectories. From these, we also computed the egocentric target distance as **z**_ego_(*t*) = − **z**_target_(*t*) − **z**_RNN_(*t*). Conjunctively, we also noted the neural activations (*r*(*t*)) during these trials. We refer to these targets collectively as **o**(*t*) below and specify whenever necessary.

### Decoding setup

We trained the decoders using **r**(*t*) as inputs and **o**(*t* + Δ) as outputs, where Δ is the temporal shift used to assess whether neural activations contain information about past states or predictions about future locations. The decoders are always trained on random trajectories and evaluated on either random or characteristic trajectories, with 320 training trials and 80 test trials. We perform 3 such train-test splits for each given data point, and save two metrics: i) the median Euclidean distance (in meters) between predicted and true positions, computed across all test samples (flattened time points and trials), and ii) the *R*^2^ computed by first flattening x,y coordinates and then summing squared residuals over all test samples. The plots in Fig. 7 report the median, first, and third quartiles for a given rank *K* over both all random seeds used to train independent RNNs and the random splits. We compute the shuffled baselines by shuffling the locations within individual trials.

### Architecture and training

We trained linear decoders using a ridge regression. The regularization hyperparameter *α* was selected via 3-fold cross-validation over the 320 training trials, evaluated on a logarithmic grid of 5 values spanning [10^−3^, 10^0^]. The value of *α* minimizing the median Euclidean error on the validation fold was then used to refit the decoder on all 320 training trials, and this final decoder was evaluated on the 80 held-out test trials.

## Supplemental Figures

**Figure S1:**
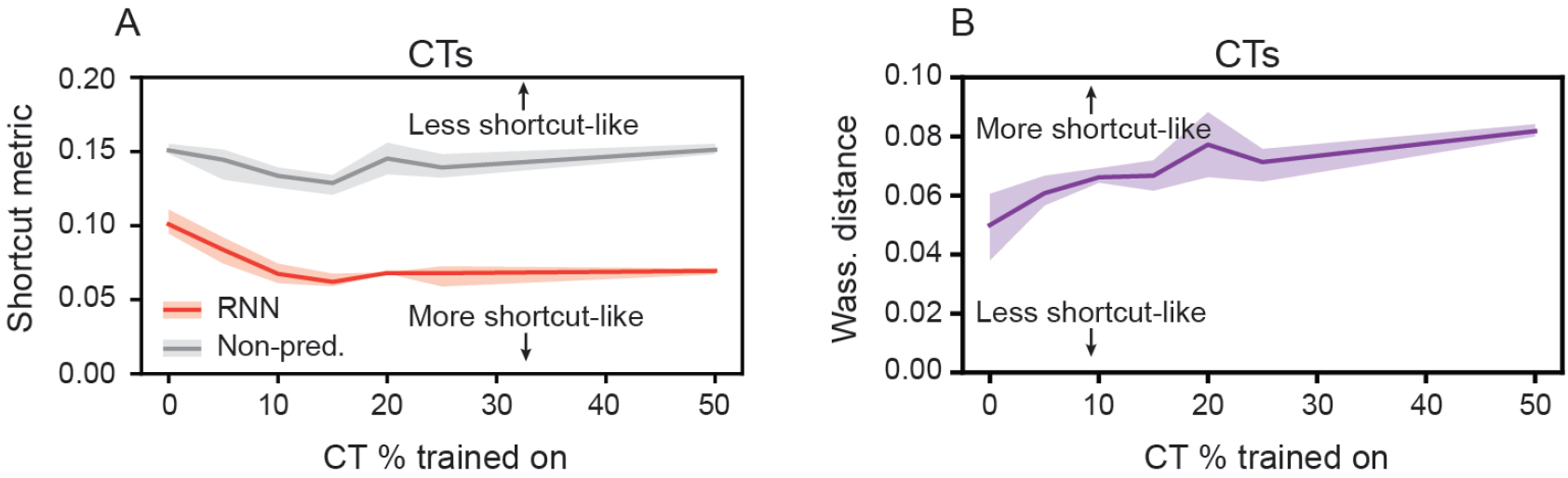
Predictive behavior of RNNs on CT trials requires with exposure to CT trials during training. (A) Shortcut metric for RNN and non-predictive models on CTs, as a function of the percent of training trials that were CTs. (B) Wasserstein distance between the distribution of RNN and non-predictive model shortcut metrics, as a function of the percent of training trials that are CTs. (A)–(B) Solid line is mean and shaded area is minimum and maximum of five independently trained networks.

**Figure S2:**
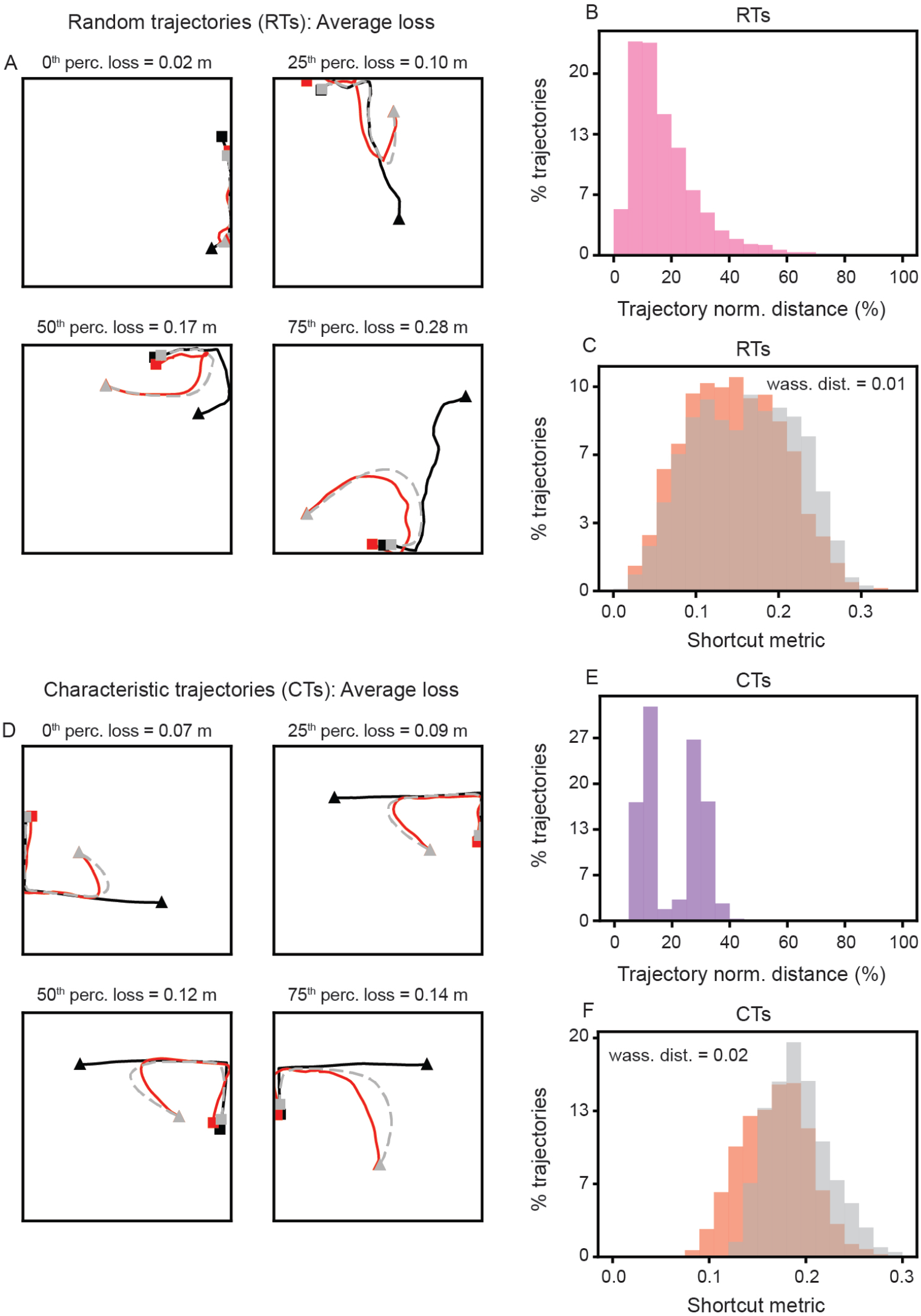
Predictivity is diminished in RNN model trained to minimize average distance to target. (A) Example RT trials for an RNN model trained to minimize the average distance to the target. Trajectories were chosen as those closest to the 0^th^, 25^th^, 50^th^, ad 75^th^ percentiles of the loss distribution, respectively. (B) Distribution of normalized distance between RNN trajectory and non-predictive model on RTs. (C) Distribution of the shortcut metric for RNN and non-predictive models on RTs. Distribution was computed across 1000 trajectories for three independently trained networks. (D)–(F) Same as (A)–(C), but for CTs. Distribution are computed across 1000 trajectories for three independently trained networks.

**Figure S3:**
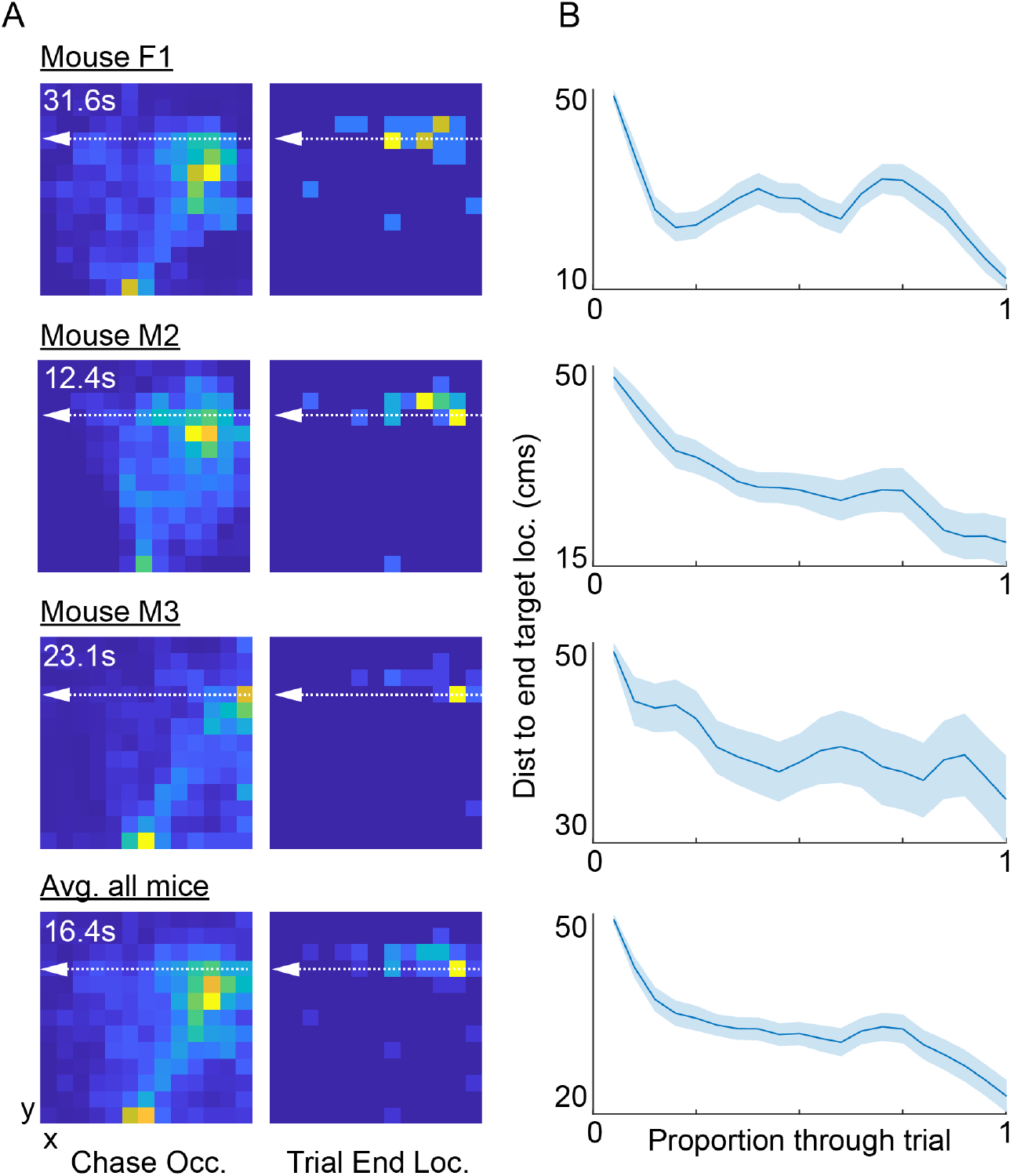
Mice learn to wait during pursuit in an environment with periodic boundaries. (A) Left column, occupancy map showing time in each arena position for 3 mice (F1, M2, M3) during periodic boundary pursuit trials. Bottom row depicts averages over all subjects. Color axis depicts occupancy in seconds (blue to yellow, zero to maximum occupancy). White numeric values in upper left corners indicate time spent in most occupied locations. Right column, occupancy map showing time in each arena position at the end of the chasing trial (including both successful and failure trials). White dashed arrow depicts trajectory of target. (B) As in Figure 5E, the average distance between the current location of the mouse vs the location of the target the end of the trial. Mice are ordered as in (A). All mice exhibited “waiting” strategies in their behavior as demonstrated by a plateau in the distance metric through the middle of the trial. Bottom row depicts average over all mice.

**Figure S4:**
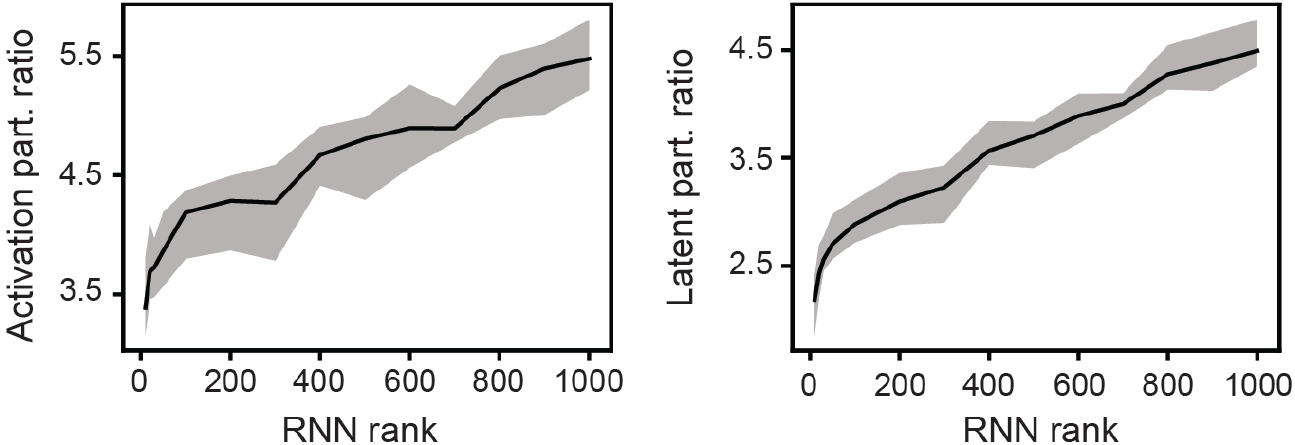
High-dimensionality of RNN activity and latent variables is also observed when using participation ratio, instead of PCA, to define dimensionality. Same as Fig. 6B, but for participation ratio instead of PCA.

**Figure S5:**
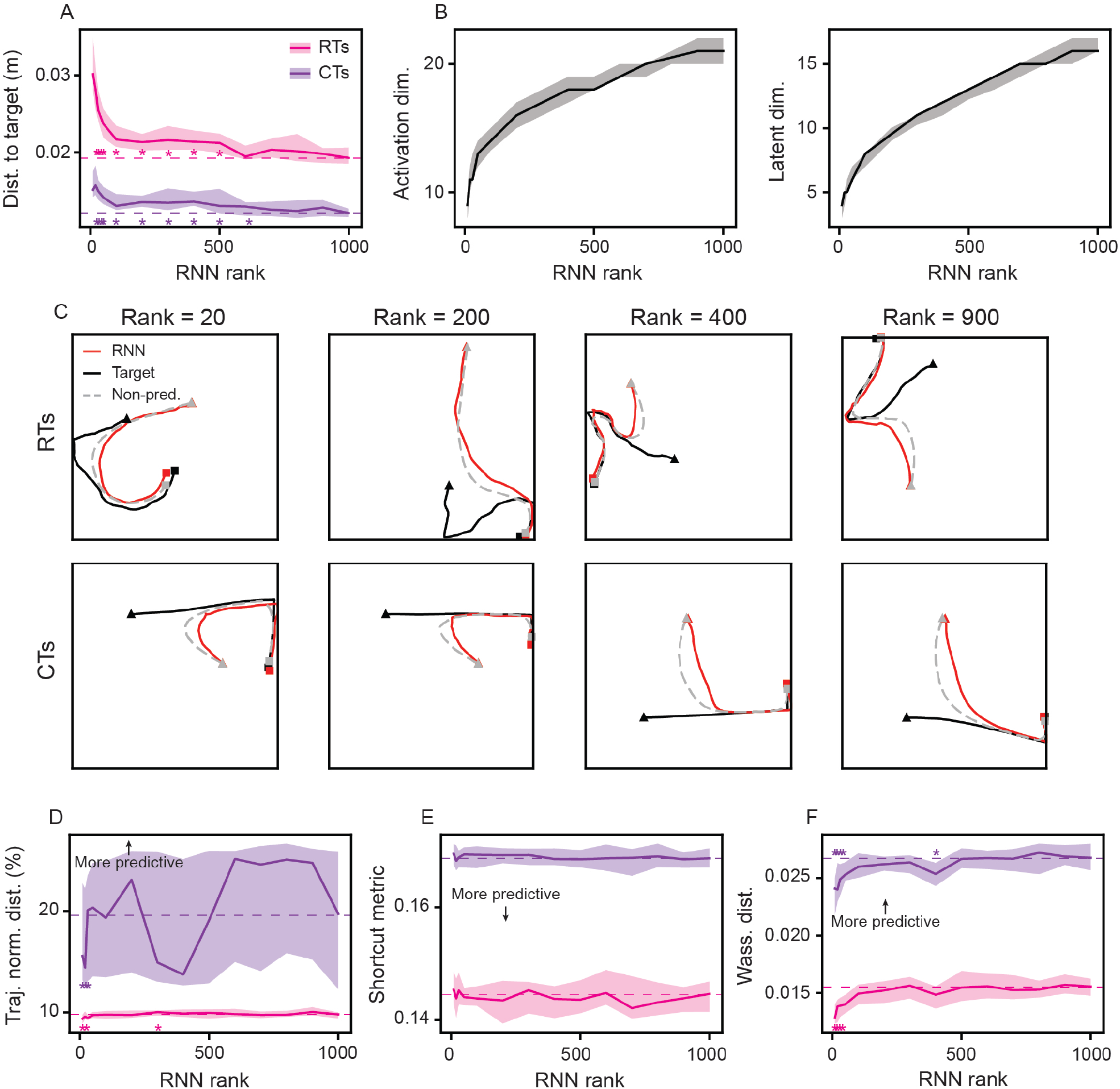
Predictivity does not emerge with increased RNN model rank, when training to minimize average distance to target. Same as Fig. 6, but for RNN whose loss function is the average distance between the target-agent and the RNN-agent, instead of the distance at the final time-point in the trial.

**Figure S6:**
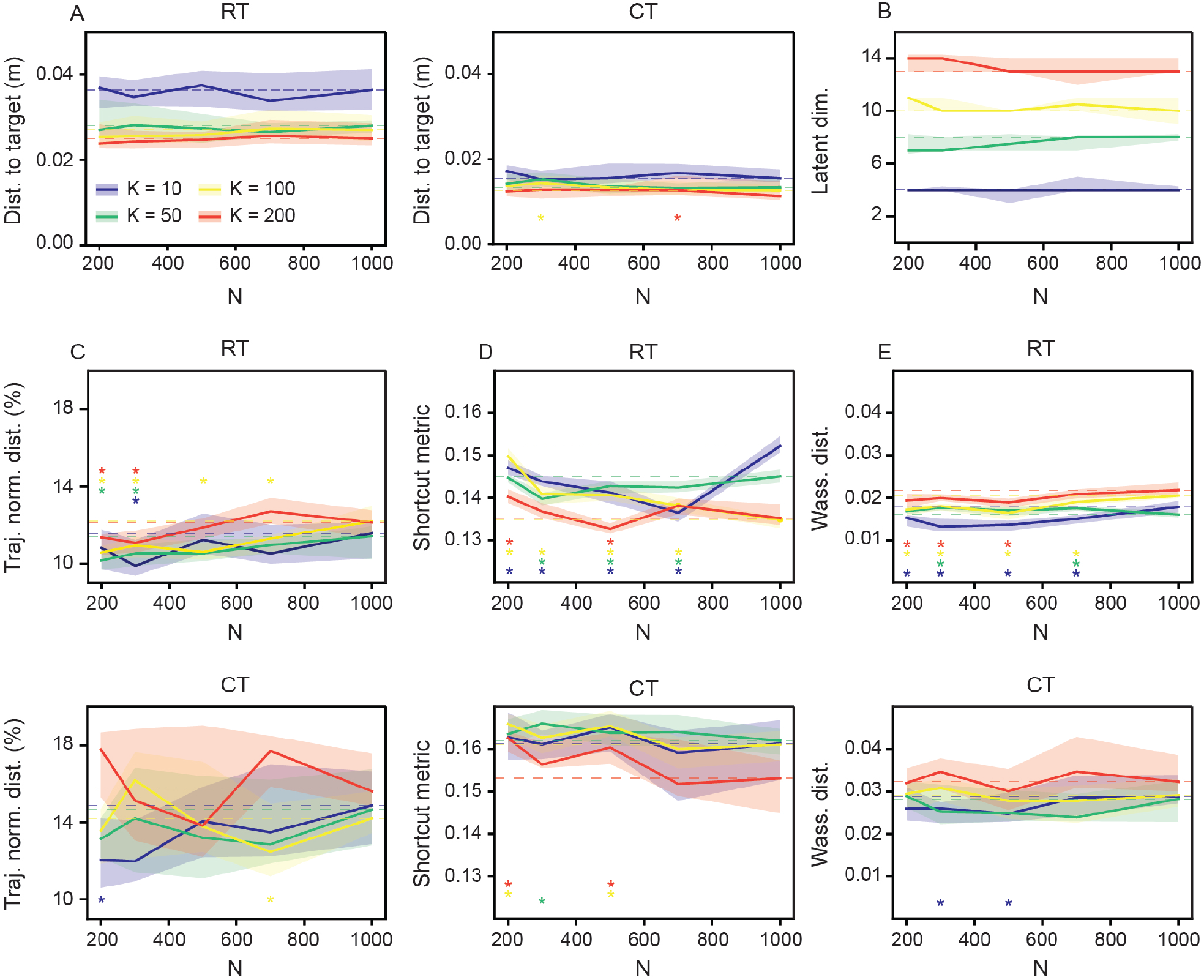
Predictivity of low rank RNNs depends on rank not network size. (A) Distance to end location of target-agent for RNNs of fixed rank and varying number of hidden units (*N* ). (B) Dimensionality (as estimated by PCA) of latent variables, as a function of *N*. (C) Normalized distance between RNN and non-predictive control model trajectories, as a function of *N* for RTs (top row) and CTs (bottom row). (D) Shortcut metric, as a function of *N*. (E) Wasserstein distance between RNN and non-predictive model shortcut metric distrubtions, as a function of *N*. (A)–(E) Solid line is median and shaded area is *±* standard deviation across 20 independently trained networks. Stars denote *N* values where two-sample Kolmogorov test showed significance with respect to *N* = 1000 model distribution (*p*−value *<* 0.05; dashed lines denote *N* = 1000 model median values).

**Figure S7:**
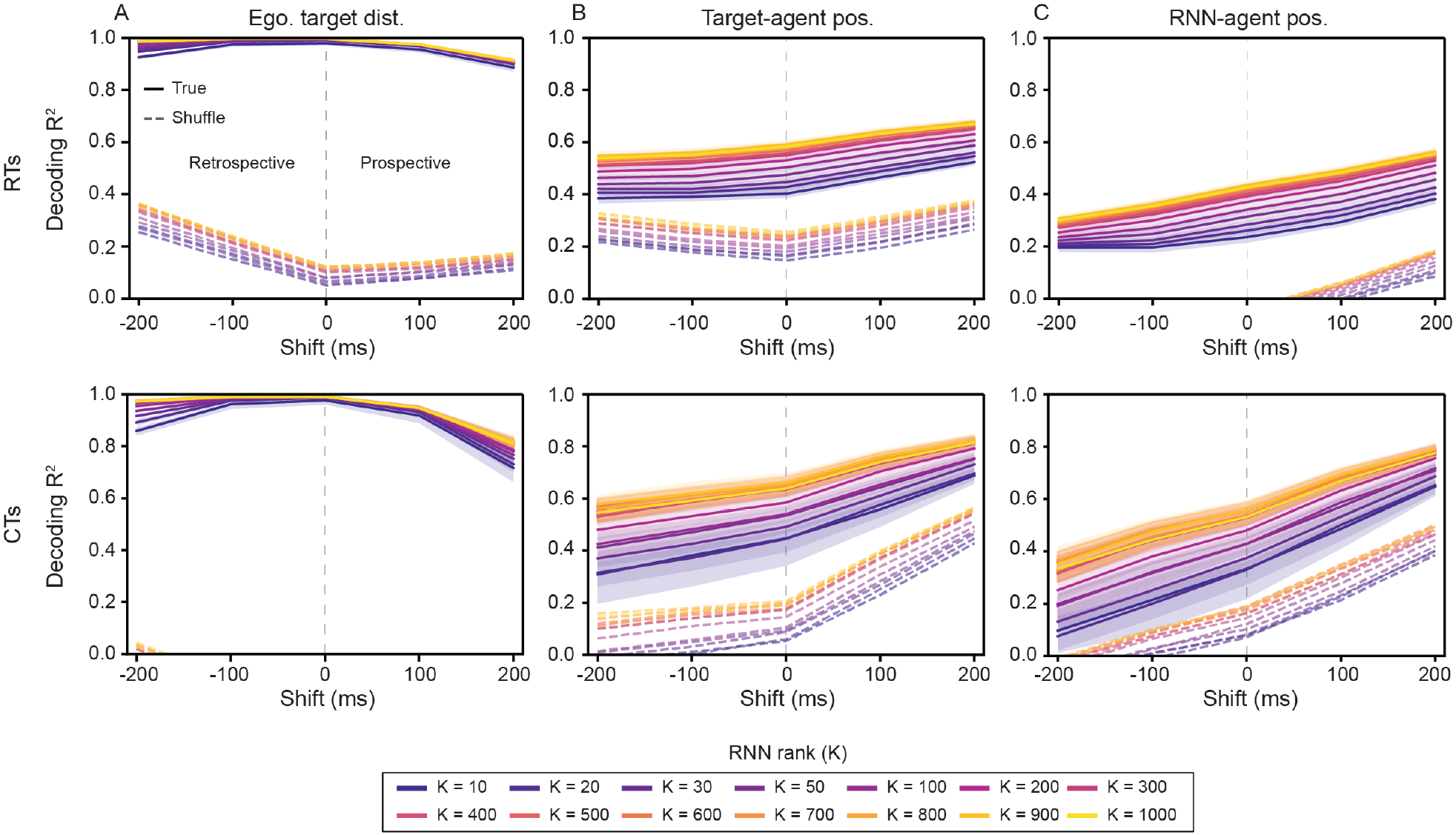
Decoding *R*^2^ of RNN-agent and target-agent position increases with increasing RNN model rank. Same as Fig. 7, but with decoding *R*^2^ instead of decoding error.

**Figure S8:**
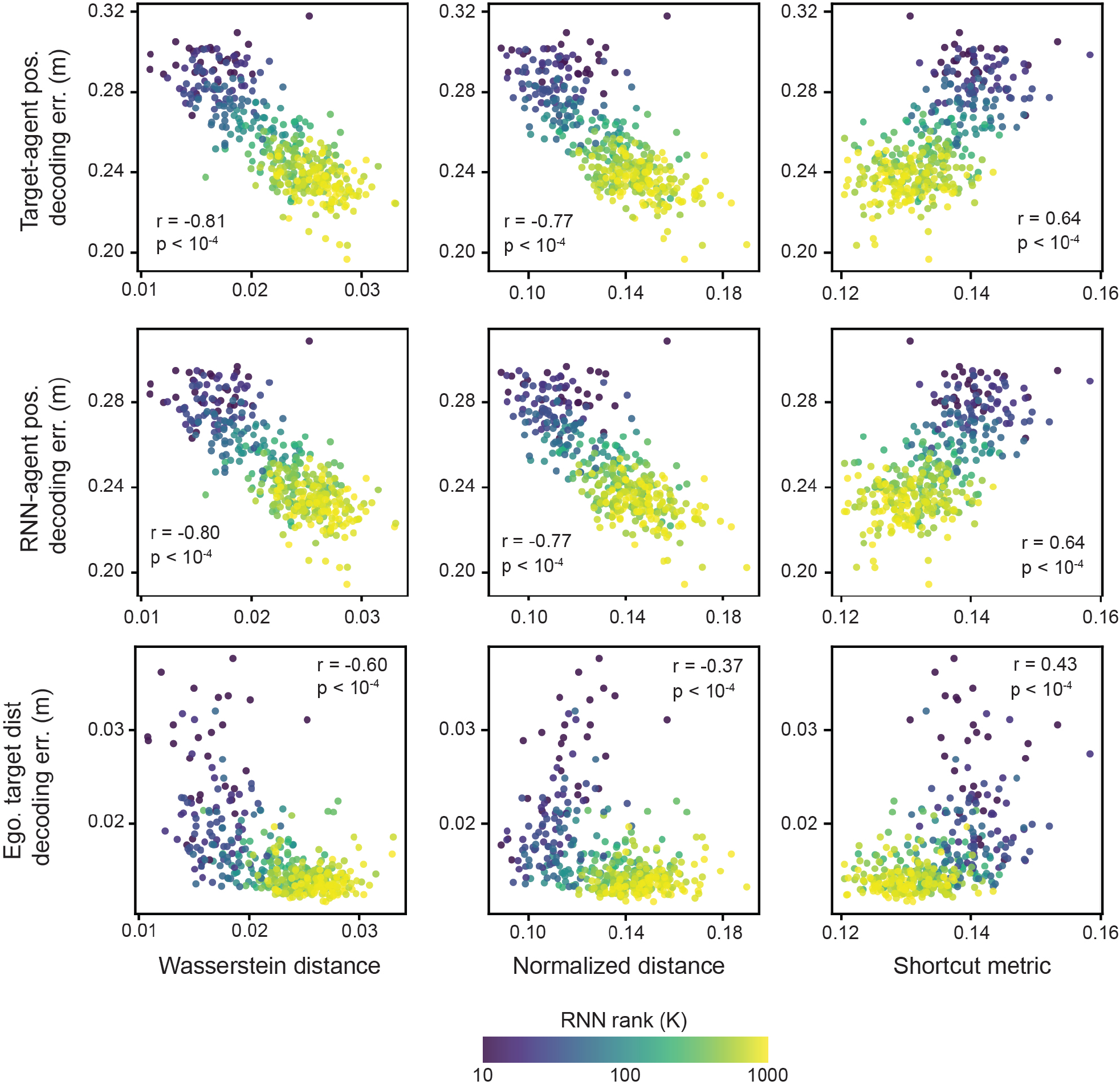
Strong correlation between predictive behavior and linear decoding error of allocentric information in RNNs. (Top row) Target-agent position decoding error as a function of Wasserstein distance, trajectory normalized distance, and shortcut metric, for all trained networks that achieved final distance to target-agent *<* 0.10 m (*n* = 415). (Middle row) RNN-agent position decoding error as a function of Wasserstein distance, trajectory normalized distance, and shortcut metric. (Bottom row) Egocentric target distance decoding error as a function of Wasserstein distance, trajectory normalized distance, and shortcut metric. Linear regression *r* and *p* − values reported in each subplot. Color of each dot denotes rank of corresponding RNN.

**Figure S9:**
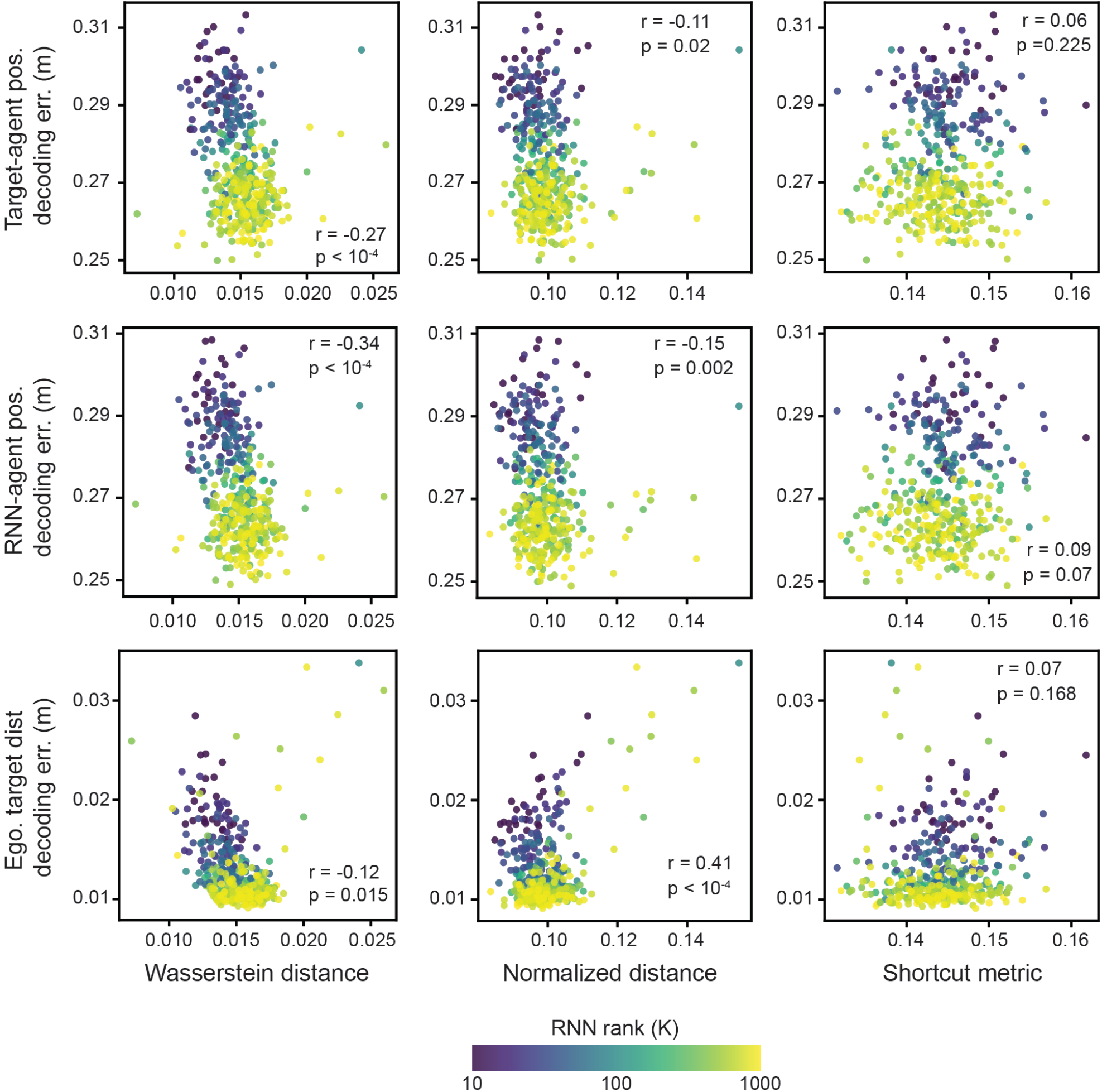
Weak correlation between predictive behavior and linear decoding error of allocentric and egocentric information in RNNs trained to minimize average distance. Same as Fig. S8, but for RNN whose loss function is the average distance between the target-agent and the RNN-agent, instead of the distance at the final time-point in the trial. *n* = 411 networks are plotted.

